# Megakaryocyte derived immune-stimulating cells regulate host-defense immunity against bacterial pathogens

**DOI:** 10.1101/2021.06.09.447810

**Authors:** Jin Wang, Jiayi Xie, Daosong Wang, Xue Han, Minqi Chen, Guojun Shi, Linjia Jiang, Meng Zhao

## Abstract

Megakaryocytes (MKs) continuously produce platelets to support hemostasis and form a niche for hematopoietic stem cell maintenance in the bone marrow. MKs are also involved in inflammation responses; however, the mechanism remains poorly understood. Here, using single-cell sequencing we identified an MK-derived immune-stimulating cell (MDIC) population exhibiting both MK-specific and immune characteristics, which highly expresses CXCR4 and immune response genes to participate in host-protective response against bacteria. MDICs interact with myeloid cells to promote their migration and stimulate the bacterial phagocytosis of macrophages and neutrophils by producing TNFα and IL-6. CXCR4^high^ MDICs egress circulation and infiltrate into the spleen, liver, and lung upon bacterial infection. Ablation of MKs suppresses the innate immune response and T cell activation to impair the anti-bacterial effects in mice under the Listeria monocytogenes challenge. Using hematopoietic stem/progenitor cell lineage-tracing mouse line, we show that MDICs are generated from infection-induced emergency megakaryopoiesis in response to bacterial infection. Overall, we identify MDICs as an MK subpopulation, which regulates host-defense immune response against bacterial infection.

## Introduction

Megakaryocytes (MKs) are large and rare hematopoietic cells in the bone marrow, which continually produce platelets to support hemostasis and thrombosis (Deutsch and Tomer, 2006). MK progenitors undergo multiple rounds of endomitosis during maturation to achieve polyploid (Chang et al., 2007; Deutsch and Tomer, 2013; Machlus and Italiano, 2013; Nagata et al., 1997; Patel et al., 2005). MKs and their progenitors also migrate between distinct microenvironments and organs for their proliferation, maturation, and biological functions (Avecilla et al., 2004; Fuentes et al., 2010; Lefrancais et al., 2017; Pal et al., 2020; Tamura et al., 2016; Wang et al., 1998). Although platelet generation is the prominent role of MKs, emerging evidence suggests that MKs have other biological functions. Mature MKs physically interact with HSCs and constitute a unique niche to HSC quiescence in the bone marrow (Bruns et al., 2014; Zhao et al., 2014). MKs also interact with other niche cells, such as osteoblasts (Dominici et al., 2009; Olson et al., 2013), non- myelinating Schwann cells (Jiang et al., 2018; Yamazaki et al., 2011), and blood vessels (Avecilla et al., 2004; Sacma et al., 2019) to further influence the attraction and retention of hematopoietic stem and progenitor cells during homeostasis and stress.

MK biased hematopoietic stem cells (HSCs) induce emergency megakaryopoiesis to actively generate MKs upon acute inflammation, which can efficiently replenish the loss of platelets during inflammatory insult (Haas et al., 2015). Studies suggested that MKs might participate in immune responses independent of their platelet generation role (Cunin and Nigrovic, 2019). MKs express multiple immune receptors, such as IgG Fc receptors and toll-like receptors (TLRs), enabling them to sense inflammation directly (Cunin and Nigrovic, 2019). Mature MKs also express major histocompatibility complex (MHC) to activate antigen-specific CD8^+^ T cells and enhance CD4^+^ T cells and Th17 cell responses through stimulating antigen processing (Finkielsztein et al., 2015; Pariser et al., 2021; Zufferey et al., 2017). Furthermore, MKs release multiple cytokines and chemokines to influence immune cells. For example, MKs produce IL-1α and IL-1β to promote arthritis susceptibility in mice resistant to arthritis (Cunin et al., 2017) and produce CXCL1 and CXCL2 to promote neutrophil efflux from the bone marrow (Köhler et al., 2011). Lung MKs contribute to thrombosis (Lefrancais et al., 2017) and, more interestingly, participate in immune responses (Pariser et al., 2021), although the relationship between lung MKs and bone marrow circulating MKs (Nishimura et al., 2015) remains unexplored. Furthermore, the recent single-cell atlas shows that MKs are heterogeneous and contain subpopulations that express multiple immune genes and are involved in inflammation response (Liu et al., 2021; Pariser et al., 2021; Sun et al., 2021; Yeung et al., 2020). However, the mechanisms that MK subpopulations regulate immune cells against bacterial pathogens remain poorly understood.

Here, by combining scRNA-seq with functional assays, we identified an MK-derived immune-stimulating cell (MDIC) population, which was generated by infection-induced emergency megakaryopoiesis, and stimulated innate immunity against bacterial infection.

## Results

### Single-cell atlas identifies an immune-modulatory subpopulation of MKs

We applied droplet-based scRNA-seq with CD41^+^ forward scatter (FSC)^high^ bone marrow MKs to explore the MK heterogeneity (Figure 1A; Figure 1-figure supplement 1A, B). To enrich accurate MKs, we further performed transcriptomic profile analysis in the phenotypically enriched MKs (Yeung et al., 2020). Our scRNA-seq successfully detected 5368 high-quality cells (Figure 1-figure supplement 1B, C), in which one MK cluster (1712 cells) and six immune cell clusters (3656 cells) were annotated according to their gene profile (Figure 1-figure supplement 1D-G) and alignment with published scRNA-seq immune cell data (Almanzar et al., 2020; Hamey et al., 2021; Pariser et al., 2021; Xie et al., 2020; Yeung et al., 2020). Our annotated MKs shared a similar gene profile with reported MKs but had a distinct gene profile with immune cells, including myeloid progenitors, basophils, neutrophils, monocytes, dendritic cells, macrophages, B cells, and T cells, in an integrated scRNA-seq analysis platform (Figure 1-figure supplement 2). Therefore, we re-clustered the transcriptionally enriched 1712 MKs into five subpopulations, termed MK1 to MK5 (Figure 1B; Figure 1-figure supplement 3A, B), which were further confirmed by the integrated scRNA-seq analysis platform to rule out the potential immune cell contamination (Pariser et al., 2021; Xie et al., 2020; Yeung et al., 2020) (Figure 1-figure supplement 3C, D). However, we noticed that mature MKs with huge sizes were captured at a relatively low rate, potentially due to the limitation in current techniques in cell purification and single-cell preparation (Liu et al., 2021; Sun et al., 2021). Enriched signature genes in Gene Ontology exhibited that MK1 and MK2 highly expressed nuclear division, DNA replication and repair genes for endomitosis (Figure 1C, D). MK3 enriched blood coagulation and thrombosis genes for platelet generation (Figure 1C, D). No signature pathways were enriched in MK4. MK5 enriched cell migration and immune response genes (Figure 1C, D; Figure 1-figure supplement 4A-E), cytokine, chemokine (Figure 1E, F; Figure 1-figure supplement 4F), and genes involved in immune cell interaction (Figure 1G, Figure 1-figure supplement 5A). MK5 also expressed signature genes in recently reported inflammatory-related MKs (Cd53, Lsp1, Anxa1, Spi) (Sun et al., 2021) and immune MKs (Ccl3, Cd52, Selplg, Sell, Adam8) (Liu et al., 2021) (Figure 1- figure supplement 5B). We also noticed that MK5 highly expressed Cxcr4 than other MK subpopulations (Figure 1H, I), although most MKs express CXCR4 (Hamada et al., 1998) (Figure 1-figure supplement 6A). To confirm this, we found that CXCR4^high^ MKs expressed MK markers (Figure 1-figure supplement 6B), and were mainly polyploid cells (Figure 1-figure supplement 6C) and had platelet generation ability (Figure 1-figure supplement 6D), although they have relatively low polyploidy (Figure 1-figure supplement 6E) and smaller cell size (Figure 1-figure supplement 6F-H). Overall, using scRNA-seq, we identified a cell type (MK5) exhibiting both MK-specific and immune characteristics, which might take part in immune responses.

**Figure 1.**
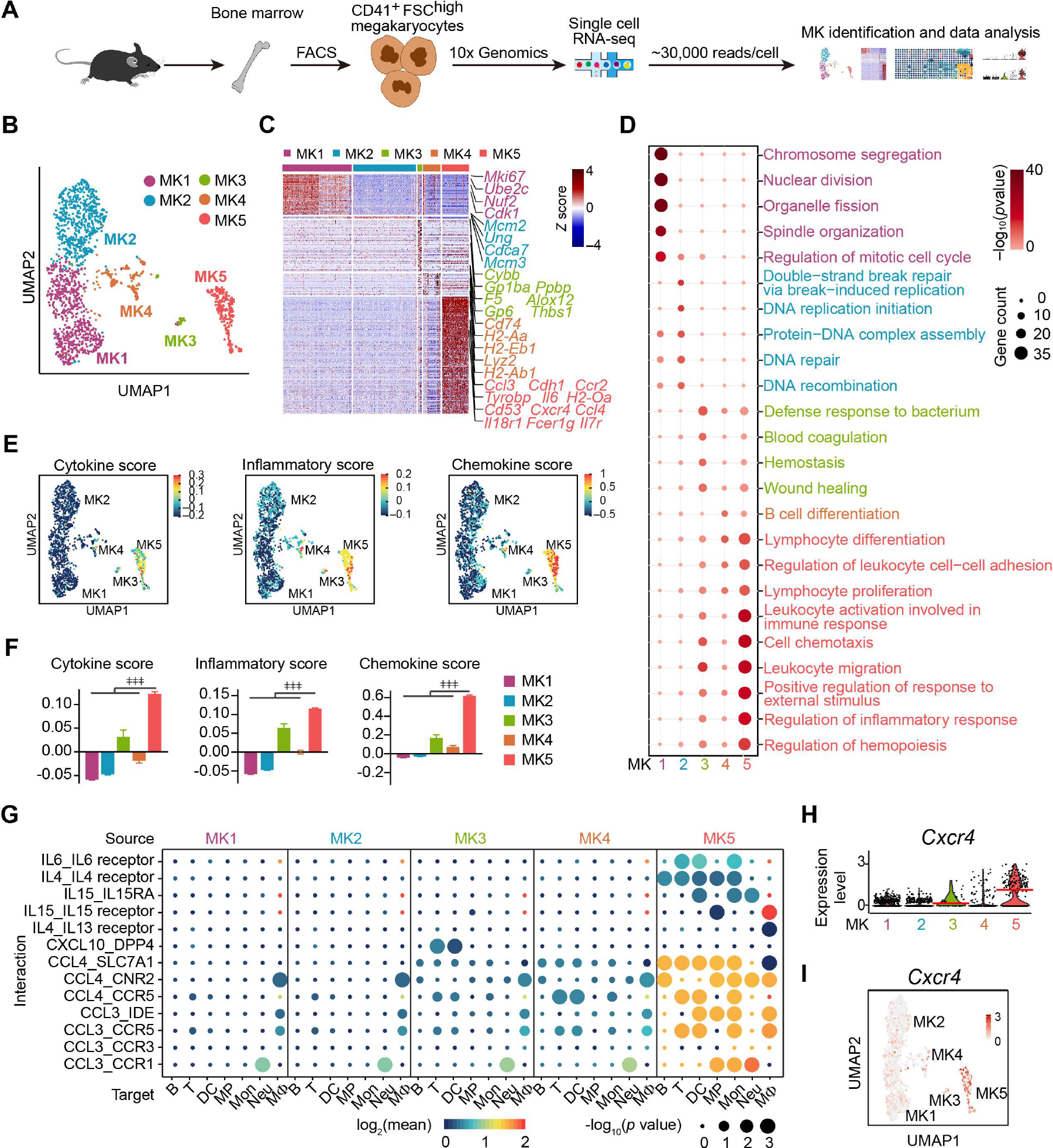
Single-cell atlas identifies an immune-modulatory subpopulation of MKs (A) Schematic strategy for MK preparation, scRNA-seq and data analysis. (B) Clustering of 1 712 bone marrow MKs. (C) Heatmap of signature gene expression in MK subpopulations (fold-change > 1.5, p value < 0.05) with exemplar genes listed on the right (top, color-coded by subpopulations). Columns denote cells; rows denote genes. Z score, row-scaled expression of the signature genes in each subpopulation. (D) Gene Ontology (GO) analysis of signature genes (fold-change > 1.5, p value < 0.05) for each MK subpopulations. GO terms selected with Benjamini–Hochberg-corrected p values < 0.05 and colored by –log10(p value). Bubble size indicates the enriched gene number of each term. (E-F) UMAP visualization (E) and statistical analysis (F) of cytokine score (left), inflammatory score (middle) and chemokine score (right) in MK1 to 5. (G) Dotplots of significant cytokine ligand (source) -receptor (target) interactions between MKs and immune cells discovered. The color indicates the means of the receptor-ligand pairs between two cell types and bubble size indicates p values. Mon, monocytes; MΦ, macrophages (Dong et al., 2020); DC, dendritic cells; Neu, neutrophils; MP, myeloid progenitors; T, T cells; B, B cells. (H-I) Violin plot (H) and feature plot (I) of selected signature genes of MK5. Red lines in (H) indicate the median gene expression. Repeated- measures one-way ANOVA followed by Dunnett’s test for multiple comparisons in (F), ǂǂ p <0.01, ǂǂǂ p <0.001. The following figure supplements are available for figure 1: Figure 1-figure supplement 1. Cell isolation, quality control and annotation of scRNA-seq data. Figure 1-figure supplement 2. Cell type identification by alignment with published scRNA- seq data. Figure 1-figure supplement 3. Identification of MK subpopulations. Figure 1-figure supplement 4. Enriched genes in MK1 to 4 and MK5. Figure 1-figure supplement 5. MK5 interacts with immune cells and express signature genes of immune MKs. Figure 1-figure supplement 6. Polyploidy, platelet generation ability and cell size of CXCR4low and CXCR4high MKs.

MDICs enhance myeloid cell mobility and bacterial phagocytosis

As MK5 enriched genes involved in myeloid cell activation (Figure 1-figure supplement 4E) and myeloid cell interactions (Figure 1G, Figure 1-figure supplement 5A), we further explored the role of MK5 in regulating myeloid immune cells. To investigate the role of MKs in regulating the innate immunity function of myeloid cells against pathogens, we challenged mice with Listeria (L.) monocytogenes, a Gram-positive facultative intracellular bacterium (Bishop and Hinrichs, 1987; Edelson and Unanue, 2000), which induce myelopoiesis (Eash et al., 2009) (Figure 2-figure supplement 1A, B). Interestingly, we noticed that more myeloid cells were associated with CXCR4^high^ MKs than with CXCR4^low^ MKs in the bone marrow of mice three days after L. monocytogenes infection (Figure 2A, B), a dramatic rise compared to randomly placed myeloid cells to MKs (Figure 2B). In supporting this, the myeloid cell-CXCR4^high^ MK association (mean distance 15.36 μm) was significantly closer than the myeloid cell-CXCR4^low^ MKs association (Figure 2C; mean distance 25.62 μm, p = 7.0×10^−4^ by KS test) in the bone marrow of mice three days after L. monocytogenes infection. Furthermore, the observed mean distance of myeloid cells to CXCR4^high^ MKs (15.36 μm) is significantly closer than the randomly placed myeloid cells to CXCR4^high^ MKs [35.37 μm, p (μ<15.36) = 1.8×10^-10^], whereas the observed mean distance of myeloid cells to CXCR4^low^ MKs (25.62 μm) is not different from random simulations [27.76 μm, p (μ<25.62) = 0.14] (Figure 2C). We also noticed that bone marrow myeloid cells were preferably adjacent to the CXCR4^high^ MK-blood vessel intersection in mice three days after L. monocytogenes infection (Figure 2-figure supplement 1C, D). These observations indicated that CXCR4^high^ MKs might regulate myeloid cells upon bacterial infection.

**Figure 2.**
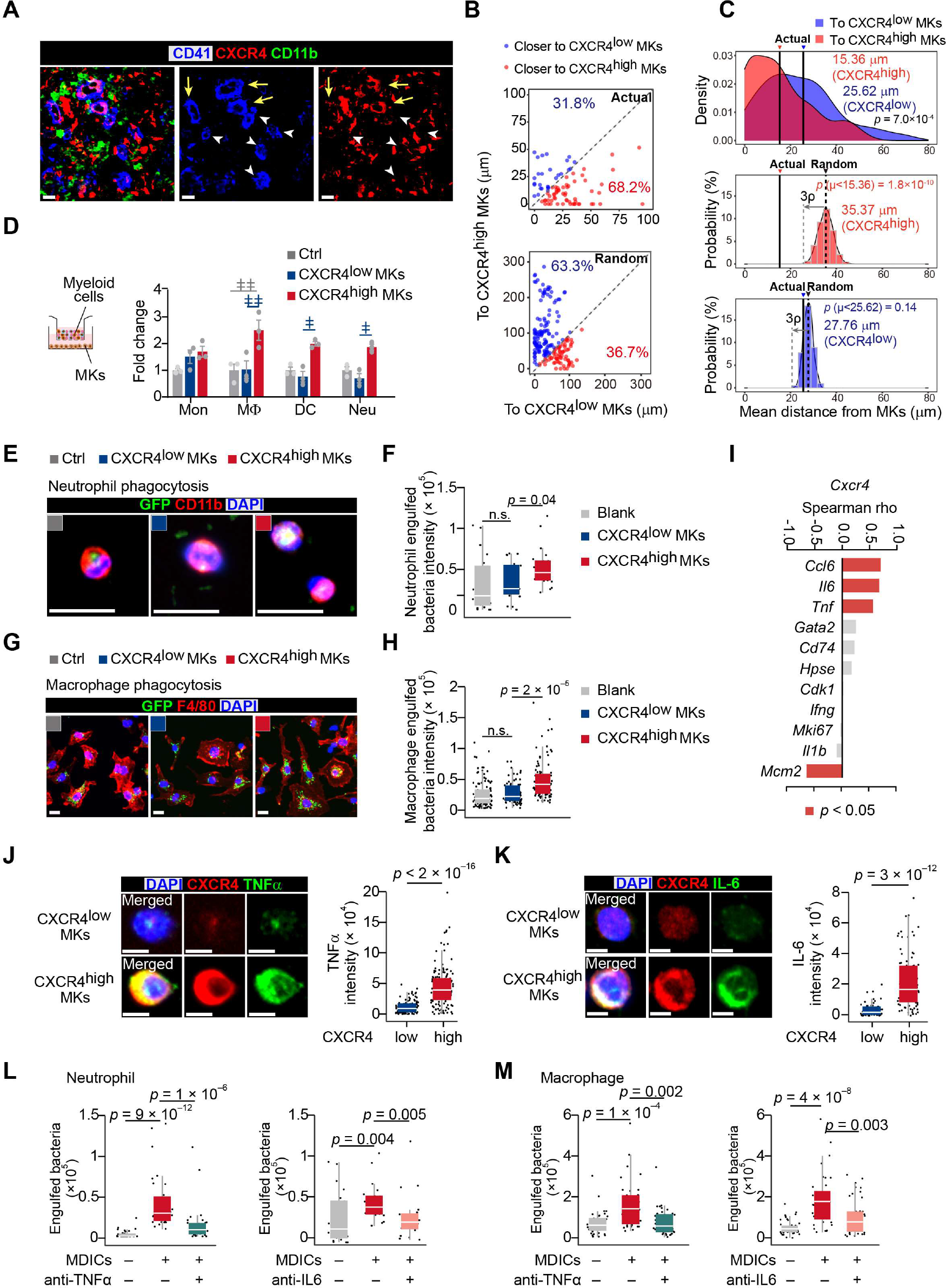
MDICs enhance myeloid cell mobility and bacterial phagocytosis (A) Distribution of myeloid cells to CXCR4^low^ or CXCR4^high^ MKs three days after L. monocytogenes infection. Representative images of MKs (blue), CXCR4 (red), and myeloid cells (green) in mouse bone marrow. Yellow arrows indicate CXCR4^high^ MKs and white arrowheads indicate CXCR4^low^ MKs. (B-C) Distance (B) and mean distance (C) of actual or randomly positioned myeloid cells to the closest CXCR4^low^ and CXR4^high^ MKs. (D) Numbers of transmigrated myeloid cells normalized to Ctrl (without MKs in the lower chambers) as indicated by transwell assays. Mon, monocytes; MΦ, macrophages; DC, dendritic cells; Neu, neutrophils. (E-F) Representative images (E) and quantification (F) of neutrophil phagocytosis capacity with or without MK co-culture as indicated. CD11b, red; E. coli, green; DAPI, blue. Blank, neutrophil without MK co-culture (Blank, n = 20; CXCR4^low^ MKs, n = 19; CXCR4^high^ MKs, n = 20). (G-H) Representative images (G) and quantification (H) of macrophage phagocytosis capacity with or without MK co-culture as indicated. F4/80, red; E. coli, green; DAPI, blue. Blank, macrophage without MK co-culture (Blank, n = 110; CXCR4^low^ MKs, n = 69; CXCR4^high^ MKs, n = 94). (I) Spearman correlation analysis between expression profiles of Cxcr4 and feature genes in MK subpopulations. (J-K) TNFα (J) and IL-6 (K) protein levels in purified CXCR4^low^ MKs and CXCR4^high^ MKs by immunostaining. (L-M) Quantification of neutrophil (L) and macrophage (M) phagocytosis with or without CXCR4^high^ MK co-culture in the absence or presence of anti-TNFα or anti-IL-6 neutralizing antibodies. Scale bars, 20 μm (A, E, G) or 5 μm (J, K). Data represent mean ± s.e.m (D) and means ± first and third quartiles (F, H, J-M). Repeated-measures one-way ANOVA followed by Dunnett’s test for multiple comparisons in (D), ǂ p <0.05, ǂǂ p <0.01. A two-sample KS test was performed to assess statistically significant (C, F, H, J-M), n.s., not significant. The following figure supplements are available for figure 2: Figure 2-figure supplement 1. L. monocytogenes promote myelopoiesis and the association of myeloid cells and the CXCR4^high^ MK-blood vessel intersection. Figure 2-figure supplement 2. TNFα and IL-6 expression in CXCR4^low^ and CXCR4^high^ MKs.

To explore how CXCR4^high^ MKs regulate myeloid cells, we interestingly found that CXCR4^high^ MKs more effectively promoted myeloid cell mobilization than CXCR4^low^ MKs in our transwell assays (Figure 2D). Furthermore, we asked whether CXCR4^high^ MKs regulate myeloid cell function against pathogens. To this aim, we incubated purified CXCR4^low^ MKs and CXCR4^high^ MKs with neutrophils or macrophages for bacterial phagocytosis analysis. We found that CXCR4^high^ MKs more efficiently enhanced the bacterial phagocytosis of neutrophils and macrophages than CXCR4^low^ MKs (Figure 2E- H). Overall, our data show that CXCR4^high^ might be a potential marker to identify a functional immune-modulating MK subpopulation. We, therefore, referred CXCR4^high^ MKs as MK-derived immune-stimulating cells (MDICs).

Our scRNA-seq also exhibited that the high expression of Cxcr4 was positively correlated with immune cell-stimulating cytokines, such as Ccl6, Tnf, and Il6 (Li et al., 2018; Rothe et al., 1993; Shapouri-Moghaddam et al., 2018) in MKs (Figure 2I). In line with this, CXCR4^high^ MKs had higher TNFα and IL-6 protein levels than CXCR4^low^ MKs (Figure 2J, K; Figure 2-figure supplement 2). The TNFα and IL-6 levels in CXCR4^high^ MKs were comparable to macrophages from mice three days after L. monocytogenes infection (Figure 2-figure supplement 2B) which are known as the primary cellular source of TNFα and IL-6 upon infection (Shapouri-Moghaddam et al., 2018). These observations suggested that MDICs might stimulate myeloid cell phagocytosis by producing TNFα and IL-6. Indeed, anti-TNFα and anti-IL-6 blocking antibodies significantly compromised the role of MDICs in stimulating bacterial phagocytosis of neutrophils and macrophages (Figure 2L, M).

### MKs stimulate both innate and adaptive immunity against bacterial pathogens

To explore the in vivo role of MKs upon L. monocytogenes infection in mice, we employed Pf4-cre; iDTR mice, in which MKs were rendered sensitive to diphtheria toxin (DT) (Zhao et al., 2014) (Figure 3A, B). Notably, MK ablation dramatically increased the bacterial burdens in the liver and spleen three days after L. monocytogenes infection (Figure 3C). We also found that MK ablation reduced the number of myeloid cells, including monocytes, macrophages, dendritic cells (DCs), and neutrophils, in the liver and spleen (Figure 3D, E), suggesting the role of MKs in promoting myeloid cells against pathogens. We further investigated how MKs regulate adaptative immunity against pathogen infection. To this aim, we challenged Pf4-cre; iDTR mice with ovalbumin (OVA)-expressing recombinant microbe (L. monocytogenes-OVA). Seven days after L. monocytogenes-OVA infection, splenocytes from control or MK ablated mice were re-stimulated with OVA peptide in vitro to assess OVA-specific T cell activation (Figure 3F). Notably, MK ablation dramatically reduced the number of CD4^+^ IFNγ^+^ Th1, CD4^+^ IL4^+^ Th2, and CD8^+^ cytotoxic T lymphocytes but did not impact the total number of CD4^+^ T cells and CD8^+^ T cells (Figure 3G). These observations demonstrated that MKs regulate innate and adaptive immunity against L. monocytogenes infection. To explore whether MDICs contribute to the immune response against bacterial pathogens, we infused the purified MDICs (CXCR4^high^ MKs) and CXCR4^low^ MKs into MK ablation mice during L. monocytogenes infection. Notably, we found that the infusion with MDICs but not with CXCR4^low^ MKs partially rescued the bacterial clearance defect in MK ablation mice (Figure 3H, I).

**Figure 3.**
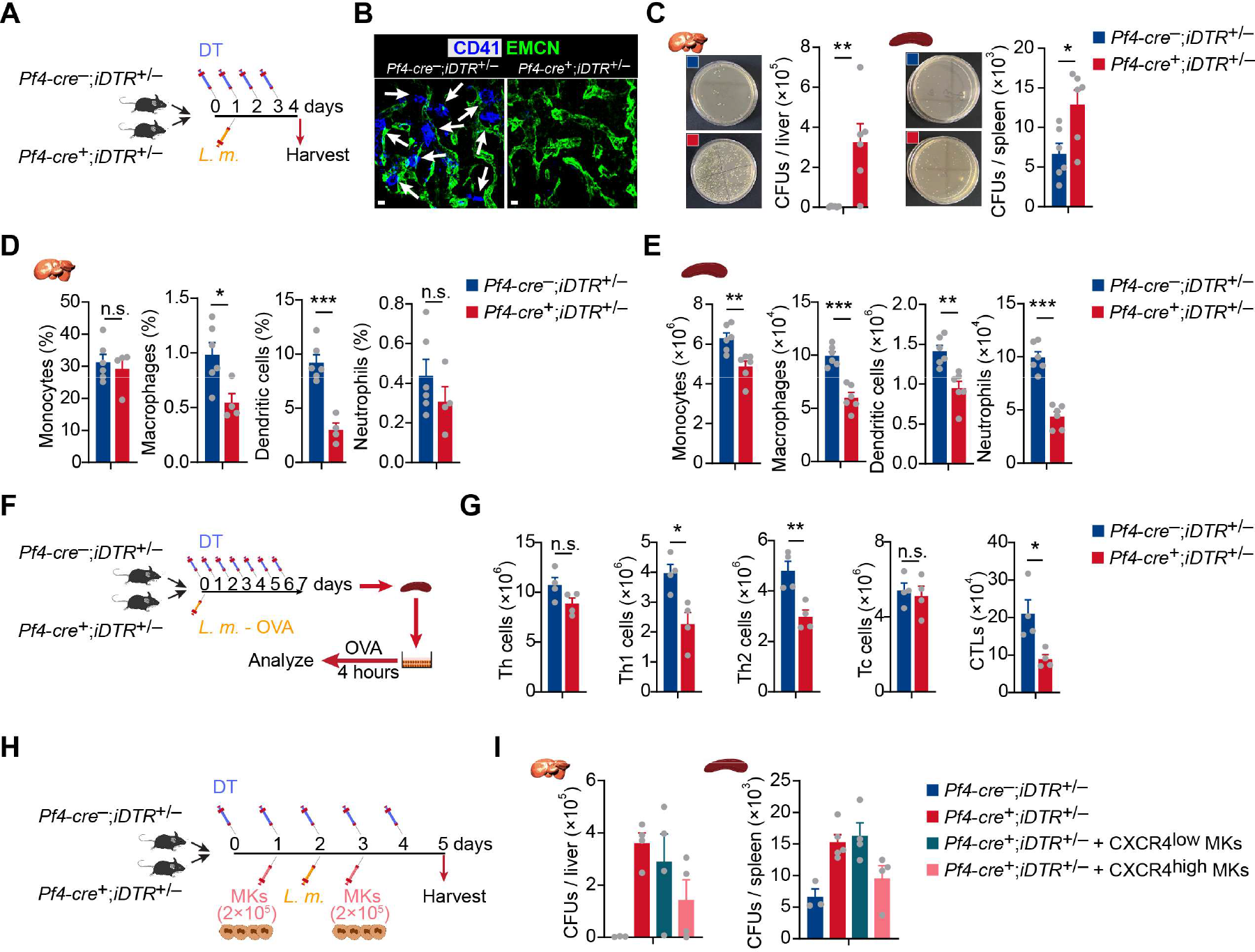
MKs stimulate both innate and adaptive immunity against bacterial pathogens (A) Schema for diphtheria toxin (DT) and L. monocytogenes administration used for the experiments shown in (B-E). (B) Representative images of MKs (blue, indicated by arrows) and vascular endothelial cells (green) in the bone marrow of mice after four daily DT treatments. (C) Bacterial burdens in the liver and spleen of Pf4-cre; iDTR mice three days after L. monocytogenes (L.m.) infection with four-time DT injections. (D-E) Myeloid cells in the liver (D) and spleen (D) of Pf4-cre; iDTR mice three days after L. monocytogenes (L.m.) infection with four-time DT injections. (F) Schema for antigen-specific T cell activation assay shown in (G). (G) Splenocytes from control or MK ablated mice seven days after L. monocytogenes-OVA257-264 infection and seven DT injections were stimulated with OVA peptide in vitro for 4 hours, and antigen-specific activated T cells were quantified (n = 4 mice). L.m.-OVA, L. monocytogenes-OVA257-264. (H) Schema for DT, L. monocytogenes administration, MK transfusing and bacterial burden determination shown in (I). (G) Bacterial burdens in the liver and spleen of Pf4-cre; iDTR mice without or with CXCR4^low^ or CXCR4^high^ MK transfused. Data represent mean ± s.e.m. Two-tailed Student’s t-test was performed to assess statistical significance, * p <0.05, ** p <0.01, *** p <0.001, n.s., not significant.

### Bacterial infection stimulates MDIC migration

High Cxcr4 expression indicated that MDICs might migrate between bone marrow microenvironment and circulation in response to infection (Suraneni et al., 2018). In line with this, our spatial distribution analysis showed that ∼80% of MKs directly contacted blood vessels three days after L. monocytogenes infection, which was much higher than in control mice (∼40%) (Figure 4A, B; Figure 4-figure supplement 1A). Furthermore, more CXCR4^high^ MKs, with small cell sizes (Figure 1-figure supplement 6F-H), were more tightly associated with blood vessels and trapped in the sinusoid than CXCR4^low^ MKs three days after L. monocytogenes infection (Figure 4C, D). However, L. monocytogenes infection did not impact the association between MKs and HSCs (Figure 4A, E), albeit the critical role of perivascular MKs in maintaining HSC quiescence (Bruns et al., 2014; Itkin et al., 2016; Zhao et al., 2014) and the dramatic HSC activation upon infection (Figure 4- figure supplement 1B).

**Figure 4.**
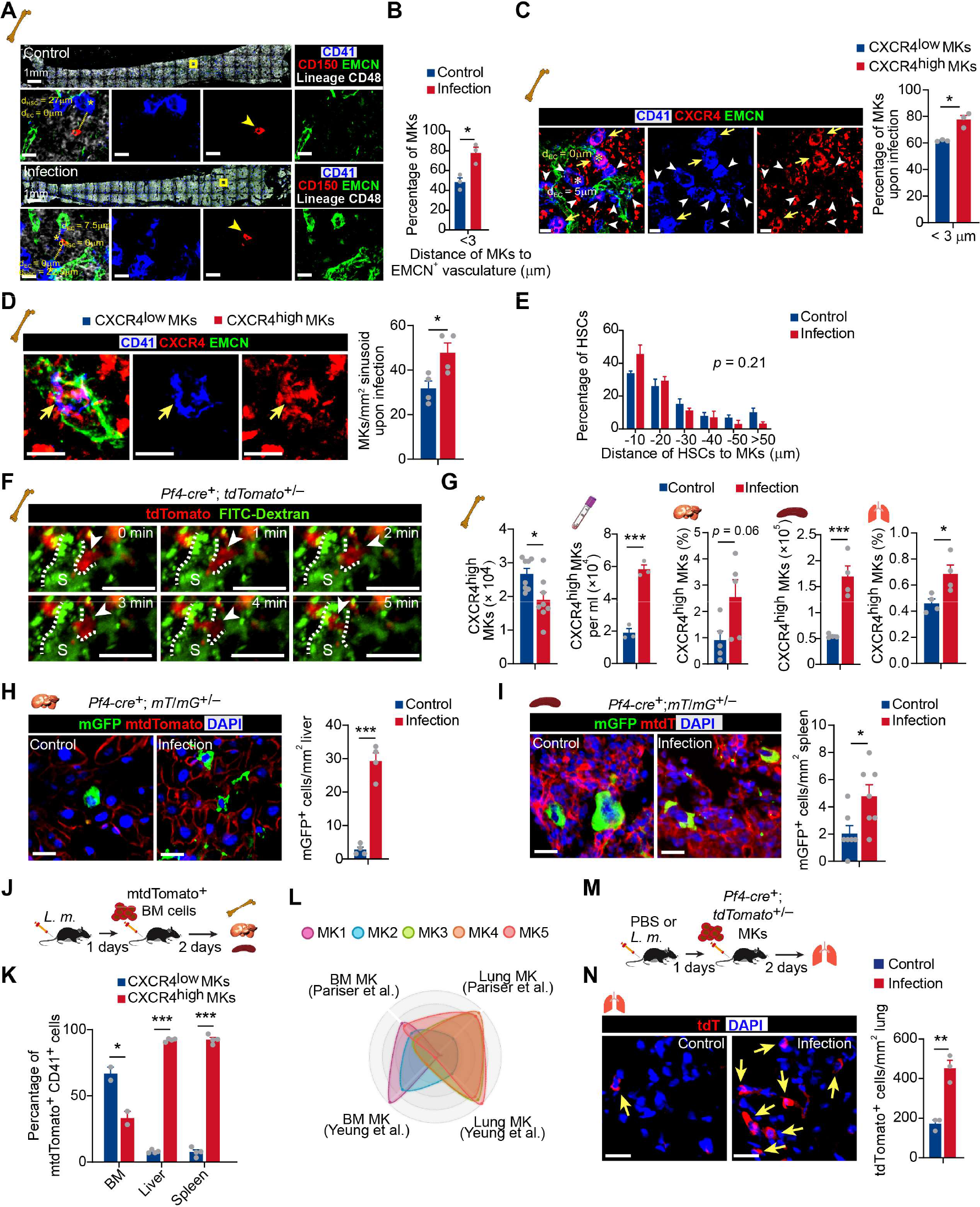
Bacterial infection stimulates MDIC migration (A) Representative image of CD41 (blue), CD150 (red), EMCN (green) and lineage cells (white) in bone marrow from control mice or mice at three days after L. monocytogenes infection. dHSC and dEC indicate the distance between the MK (blue, marked with an asterisk) and the closest HSC (red), endothelial cell (green), respectively. Yellow boxes indicate the locations of the magnified images. Arrowheads indicate HSCs. EMCN, endomucin; EC, endothelial cell. (B) Comparison of the distance between MKs to ECs (n = 119 control and 103 infected MKs) in the bone marrow of control mice or mice at three days after L. monocytogenes infection. (C) Comparison of the distance between CXCR4^low^ or CXCR4^high^ MKs and endothelial cells (ECs) in the bone marrow of mice three days after L. monocytogenes infection (n = 68 CXCR4^low^ and 78 CXCR4^high^ MKs). CD41 (blue), CXCR4 (red) and EMCN (green). Yellow arrows indicate CXCR4^high^ MKs, while white arrowheads indicate CXCR4^low^ MKs. (D) Representative immunofluorescent staining images (left) and quantification (right) of CXCR4 (red) labeled MKs (blue) egressed into sinusoids (green) upon infection (n = 46 CXCR4^low^ MKs and 69 CXCR4^high^ MKs in four biological replicates). Yellow arrows indicate CXCR4^high^ MKs. (E) Comparison of the distance between HSCs to MKs (n = 96 control and 127 infected HSCs, p = 0.21 by two- sample KS test) in the bone marrow of control mice or mice at three days after L. monocytogenes infection. (F) Visualization of MK migration (red, arrowhead) into sinusoids (green) by live imaging in the bone marrow of Pf4-cre^+^; tdTomato^+/-^ mice 24 hours after L. monocytogenes infection (Movie S1). “S” indicates sinusoid and dashed lines demarcate the border of sinusoids. (G) Quantification of CXCR4^high^ MKs in bone marrow, peripheral blood, liver, spleen, and lung of control mice and mice three days after L. monocytogenes infection. (H-I) Representative images and quantification of mGFP^+^ cells (green) in the liver (H) and spleen (I) from control mice or mice three days after L. monocytogenes infection (n = 4 and 7 mice, respectively). (J) Schema of mtdTomato^+^ bone marrow (from R26R^mT/mG^ mice) cell perfusion in control and L. monocytogenes infected recipients. (K) The percentage of CXCR4^high^ mtdTomato^+^ MKs and CXCR4^low^ mtdTomato^+^ MKs in bone marrow, liver and spleen of control or infected recipients were analyzed two days after mtdTomato^+^ bone marrow cells were perfused. (L) Radar chart showing transcriptomic similarities of bone marrow MK subpopulations with reported BM and lung MK datasets (Pariser et al., 2021; Yeung et al., 2020). (M) Schema for transfer experiments using tdTomato^+^ MKs from Pf4-cre^+^; tdTomato^+/-^ mice into control recipients or recipients one day following L. monocytogenes infection. (N) Representative images (left) and quantification (right) of tdTomato^+^ MKs in the lung of control or infected recipients two days after cell perfusion (n = 3 mice). Arrows indicate tdTomato^+^ MKs in the lung. Scale bars without indicated, 20 μm. A two-sample KS test was performed to assess statistical significance in (E). Two-tailed Student’s t-test was performed to assess statistical significance except (E), * p <0.05, ** p <0.01, *** p <0.001, n.s., not significant. The following figure supplements are available for figure 4: Figure 4-figure supplement 1. The effects of association of MKs and blood vessels, HSC activation, and MK numbers in bone marrow upon bacterial infection. Figure 4-figure supplement 2. scRNA-seq of MKs from mice upon bacterial infection. Figure 4-figure supplement 3. Cxcl12 expression upon bacterial infection. Figure 4-figure supplement 4. Cell cycle and apoptosis of CXCR4high MKs, and CXCR4low MK numbers in different organs upon bacterial infection. Figure 4-figure supplement 5. Immune gene expression in bone marrow and lung MKs.

To further explore the dynamic migration of MKs upon pathogen infection, we developed a live imaging method to trace MK migration in the bone marrow (seen in the method). Using Pf4-cre; tdTomato mice and in vivo live imaging approach, we observed that small tdTomato^+^ MKs rapidly migrated into sinusoids without rupture or platelet release upon infection (Figure 4F, Figure 4-video 1). In contrast, MKs with large sizes showed much slower migration (Figure 4-video 1). Additionally, MDICs decreased in the bone marrow three days after L. monocytogenes infection but with a similar proliferation and apoptosis rate compared to CXCR4^low^ MKs (Figure 4G, Figure 4-figure supplement 1C-F), indicating MDICs might migrate out of bone marrow. Consistent with this, the frequency of MK5, which enriched MDICs, decreased in bone marrow after L. monocytogenes infection in our single-cell atlas (Figure 4-figure supplement 2). Furthermore, we found that L. monocytogenes infection decreased the expression of CXCL12, the ligand of CXCR4 (Sugiyama et al., 2006), in bone marrow but increased CXCL12 expression in the lung, liver, and spleen (Figure 4-figure supplement 3), indicating that MDICs might migrate from bone marrow to other tissues. In line with this, MDICs increased in the peripheral blood and organs, including the liver, spleen, and lung three days after L. monocytogenes infection without an alternation of cell cycle and apoptosis, whereas CXCR4^low^ MKs did not differ except for a slight increase in the liver (Figure 4G; Figure 4-figure supplement 4).

To further explore how MKs migrate in organs during bacterial infection in vivo, we employed Pf4-cre; Rosa26-cell membrane-localized tdTomato cell membrane-localized EGFP (R26R^mT/mG^) mice in which cell membrane-localized EGFP (mGFP) expresses exclusively in MK lineage (Tiedt et al., 2007). We found that mGFP^+^ MKs increased dramatically in the liver (10.4-fold increased) and spleen (2.33-fold increased) three days after L. monocytogenes infection (Figure 4H, I). To further confirm the tissue infiltration of MKs upon infection, we intravenously injected membrane-localized tdTomato (mtdTomato) expressing bone marrow cells from R26R^mT/mG^ mice into control recipients or recipients infected with L. monocytogenes one day before mtdTomato^+^ cell perfusion (Figure 4J). We found that two days after mtdTomato^+^ cell perfusion, engrafted mtdTomato^+^ CXCR4^high^ MDICs more efficiently infiltrated into the liver (92.1%) and spleen (92.5%); by contrast, mtdTomato^+^ CXCR4^low^ MKs (66.7%) migrated to the bone marrow (Figure 4K).

As the lung is an important site for platelet generation (Lefrancais et al., 2017), we aligned our MK sc-RNAseq data with lung MKs (Pariser et al., 2021; Yeung et al., 2020), and found that MK5, MK4, and MK3 showed similar gene profiles with lung MKs (Figure 4L). Moreover, MK5 enriched more inflammatory pathway genes, antigen processing, and presentation pathway after L. monocytogenes infection, which enabled MK5 to achieve a more similar transcriptional profile as the lung MKs than control MK5 (Figure 4-figure supplement 5). Interestingly, we found that engrafted tdTomato^+^ MKs (from Pf4-cre; tdTomato mice) more efficiently infiltrated the lungs in the infected recipients than in the control recipients (Figure 4M, N).

### Acute inflammation induced emergency megakaryopoiesis generates MDICs upon infection

Infection-induced emergency megakaryopoiesis compensates the platelet consumption (Verschoor et al., 2011). Consistently, we observed that MKs were ruptured in the bone marrow three days after L. monocytogenes infection to recover the reduced platelets post-L. monocytogenes infection (Couldwell and Machlus, 2019; Nishimura et al., 2015) (Figure 5A-C). However, MDICs were increased at 18 hours after L. monocytogenes infection and substantially declined at 72 hours in bone marrow, whereas CXCR4^low^ MKs remained unchanged upon infection (Figure 5D). As MK-committed HSCs drive infection-induced emergency megakaryopoiesis (Haas et al., 2015), we asked whether emergency megakaryopoiesis also generates MDICs to participate in the host-defense response. To this aim, we employed Scl-creER; tdTomato mice (Göthert et al., 2005) to monitor the HSPC derived emergency megakaryopoiesis upon bacterial infection. Eighteen hours after tamoxifen recombining tdTomato in HSPCs and L. monocytogenes infection (Figure 5E), we observed that tdTomato^+^ HSPCs derived tdTomato^+^ CXCR4^high^ MDICs rapidly increased in the bone marrow, similar to the platelet-generating MKs (tdTomato^+^ CXCR4^low^ MKs) (Figure 5F), without a noticeable rise of hematopoietic progenitors (Figure 5G). Overall, our observations indicated that MDICs might be generated by emergency megakaryopoiesis to stimulate pathogen defense.

**Figure 5.**
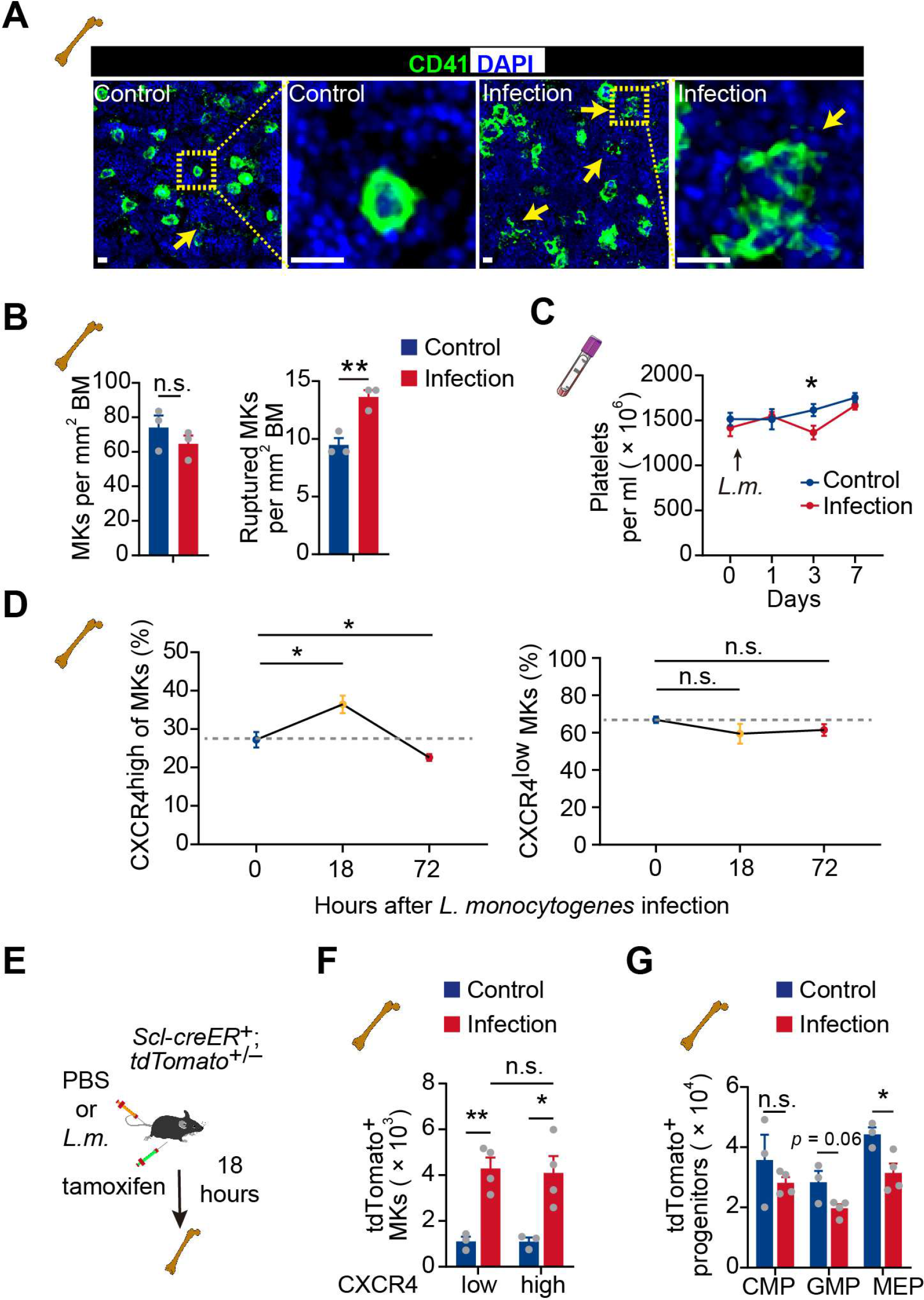
Acute inflammation induces emergency megakaryopoiesis of MDICs (A-B) Representative images (A) and statistical analysis (B) of CD41 (green) and DAPI (blue) in bone marrow from control mice or mice three days after L. monocytogenes infection. Arrows indicate ruptured MKs, yellow boxes indicate the locations of the magnified images. (C) Platelets in peripheral blood in control mice or mice after L. monocytogenes infection on indicated days. (D) The dynamics percentage of CXCR4^high^ MKs (left) or CXCR4^low^ MKs (right) in the bone marrow of L. monocytogenes-challenged mice within 72 hours of infection. (E) Schema for HSC lineage tracing upon L. monocytogenes infection using Scl-creER^+^; tdTomato^+/-^ mice. (F-G) Cell numbers of tdTomato^+^ CXCR4^low^ MKs and tdTomato^+^ CXCR4^high^ MKs (F), and tdTomato^+^ progenitors (G) in the bone marrow of control and L. monocytogenes infected Scl-creER^+^; tdTomato^+/-^ recipients 18 hours after L. monocytogenes infection and tamoxifen administration. CMP, common myeloid progenitor; GMP, granulocyte-monocyte progenitor; MEP, megakaryocyte-erythroid progenitor. Scale bars, 20 μm. Data represent mean ± s.e.m. Two-tailed Student’s t-test was performed to assess statistical significance, * p <0.05, ** p <0.01, n.s., not significant.

## Discussion

MKs express multiple immune receptors, which participate in megakaryocyte maturation, platelet activation, and potentially influence neutrophils and the adaptive immune cells (Cunin and Nigrovic, 2019). Accordingly, MKs prevent the spread of dengue virus infection by enhancing the type 1 interferons pathway in murine and clinical biospecimens (Campbell et al., 2019) and contribute to cytokine storms in severe COVID-19 patients (Bernardes et al., 2020; Ren et al., 2021; Stephenson et al., 2021). Recent scRNA-seq studies suggested the existence of MK subpopulations for inflammation responses (Liu et al., 2021; Pariser et al., 2021; Sun et al., 2021; Wang et al., 2021); however, the mechanism that MKs regulate immune response remains elusive. Here, we identified that MK5 has both MK and immune cell characteristics, producing platelets and enriching immune response genes. We also demonstrated the role of this cell type, a CXCR4^high^ MK subpopulation (MDICs) in recruiting and stimulating innate myeloid cells by producing TNFα and IL-6, for bacterial phagocytosis.

Normal HSC to MK development takes 11-12 days in humans and 4 days in mice; However, emergency megakaryopoiesis takes less than a day to generate MKs upon inflammation stress (Couldwell and Machlus, 2019; Liu et al., 2021; Sun et al., 2021) (Figure 5D). Previously, researchers believed that the emergency megakaryopoiesis mainly contributes to the replenishment of damaged platelets upon acute inflammation (Haas et al., 2015). Here, for the first time, we found that emergency megakaryopoiesis also quickly generated MDICs to facilitate immune responses against bacterial infection.

A recent report showed that the lung is a reservoir of MKs for platelet production (Lefrancais et al., 2017). Other works also indicate that lung MKs share a similar transcriptional profile with lung DCs and participate in pathogen infection (Boilard and Machlus, 2021; Pariser et al., 2021). However, the correspondence between MKs in the lung and bone marrow remains unexplored. Neonatal lung MKs lack the immune molecules in adult lung MKs (Pariser et al., 2021), which indicated that lung MKs might have distinct developmental origins. Similarly, MKs are observed to egress and migrate to the pulmonary capillary under stresses (Davis et al., 1997). Our works suggested that lung MKs might migrate from bone marrow upon infection challenges, although more detailed investigations are warranted in future studies.

## Materials and Methods

### Key resources table

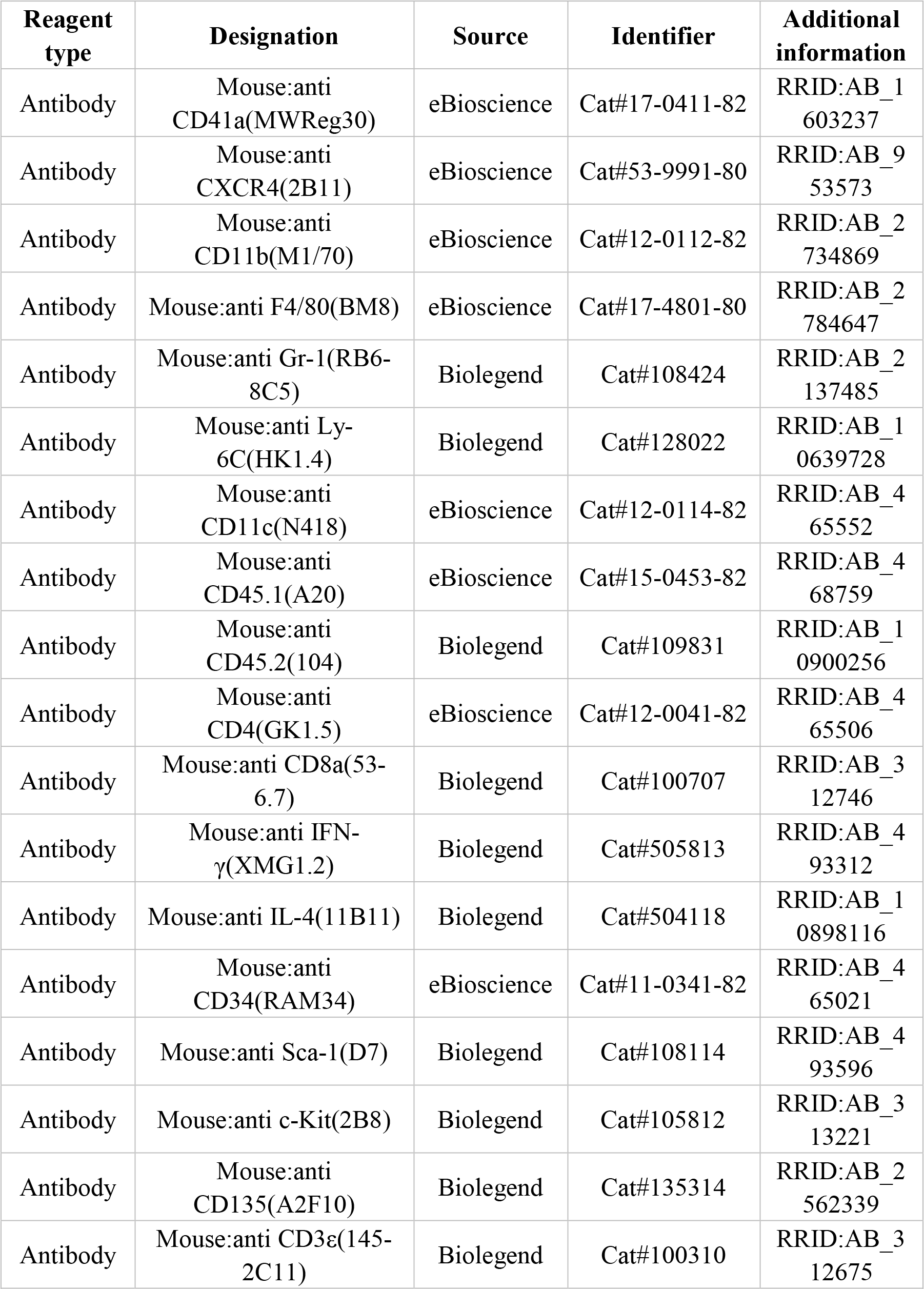
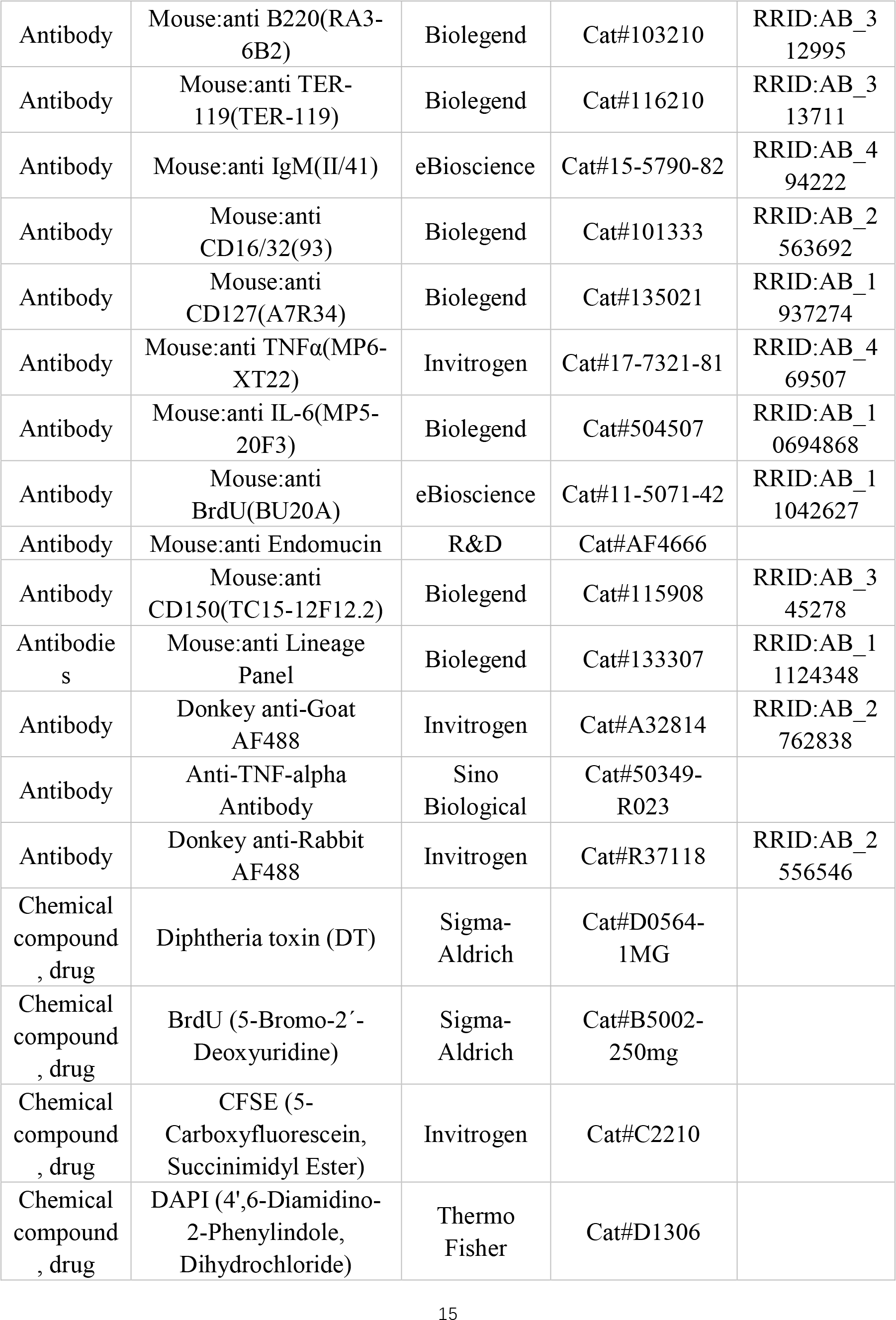
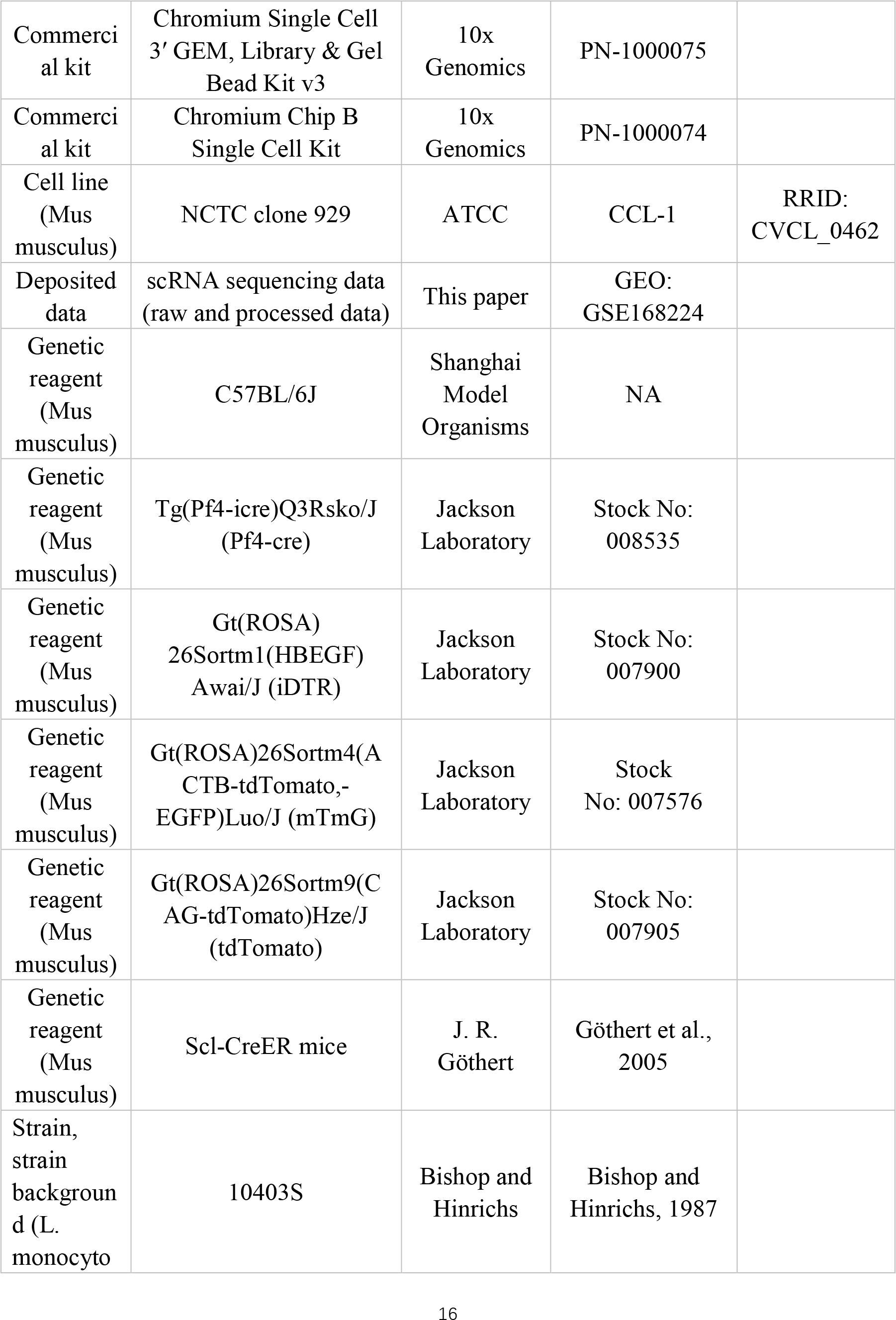
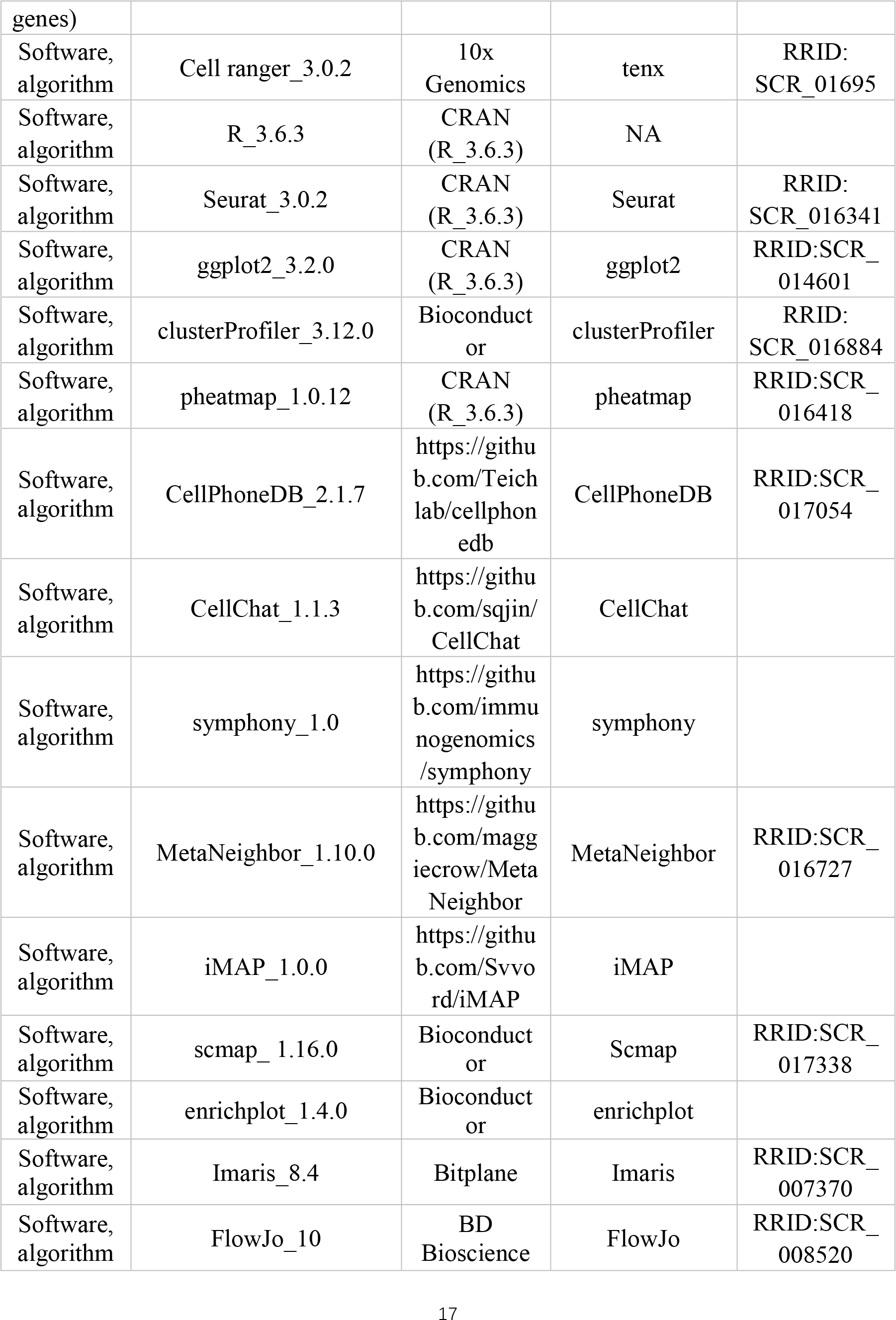
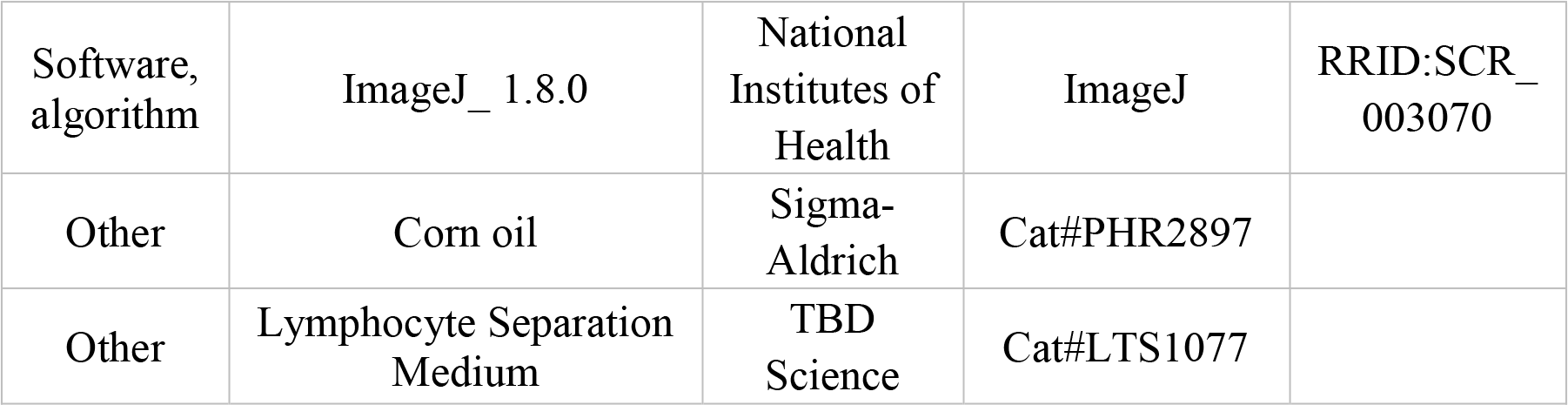

### Mice

C57BL/6-Tg(Pf4-cre)Q3Rsko/J (Pf4-cre), C57BL/6-Gt(ROSA) 26Sortm1(HBEGF) Awai/J (iDTR), C57BL/6-Gt(ROSA)26Sortm4(ACTB-tdTomato,-EGFP)Luo/J (mTmG), Gt(ROSA)26Sortm9(CAG-tdTomato)Hze (tdTomato) mice were obtained from the Jackson Laboratory. Scl-creER mice were provided by J. R. Göthert. All mice were maintained in the C57BL/6 background. Animals were blindly included in the experiments according to genotyping results as a mix of male and female. All animal experiments were performed according to protocols approved by the Sun Yat-sen University animal care and use committee.

### Bacteria and infections

Listeria (L.) monocytogenes infection was performed as described with minor modifications (Edelson and Unanue, 2000; Verschoor et al., 2011). In brief, wild type L. monocytogenes strain 10403S grown to exponential phase at 37 °C in TSB media was injected intravenously at a dose of 2 500 colony-forming units (CFUs) to determine spleen and liver bacterial burdens three days after infection. Recombinant L. monocytogenes expressing the chicken ovalbumin peptide (OVA257-264) (L.m. – OVA257-264) was injected intravenously at a dose of 2500 CFUs to determine activated spleen T cells seven days after infection. Escherichia (E.) coli wild type strain 85344 expressing GFP was constructed as previously described (Feng et al., 2020). GFP labeled E. coli was grown to exponential phase at 37 °C in LB media and washed with PBS before being suspended for phagocytosis assays.

### Antibodies for flow cytometry analysis and cell sorting

For cell sorting and analysis, monoclonal antibodies to CD41 (MWReg30, eBioscience), CXCR4 (2B11, eBioscience), CD11b (M1/70, eBioscience), F4/80 (BM8, eBioscience), Gr-1 (RB6-8C5, Biolegend), Ly6C (HK1.4, Biolegend), CD11c (N418, eBioscience), CD45.1 (A20, eBioscience), CD45.2 (104, Biolegend), CD4 (GK1.5, eBioscience), CD8 (53-6.7, Biolegend), INF-γ (XMG1.2, Biolegend), IL4 (11B11, Biolegend), CD34 (RAM34, eBioscience), Sca-1 (D7, Biolegend), c-kit (2B8, Biolegend), CD135 (A2F10, Biolegend), CD3ε (145-2C11, Biolegend), CD45R (RA3-6B2, Biolegend), TER-119 (Ter- 119, Biolegend), IgM (II/41, eBioscience), FγRII (93, Biolegend), IL-7R (A7R34, Biolegend), TNFα (MP6-XT22, Invitrogen) and IL-6 (MP5-20F3, Biolegend) were used where indicated.

### Flow cytometry and cell sorting

Bone marrow cells were isolated from mouse femora and tibiae as previously reported (Jiang et al., 2018). Splenocytes were mechanically dissociated in PBS with 2% FBS. Peripheral blood was collected from the retro-orbital sinus and anticoagulated by K2- EDTA. Those three kinds of cells then underwent red blood cell lysis for 5 min using 0.16 M ammonium chloride solution. Liver cells were mechanically dissociated and lysed using 0.16 M ammonium chloride solution, followed by gradient sedimentation using a density reagent (LTS1077, TBD Science) following the manufacturer’s instruction. Cell sorting was performed using a cell sorter (MoFlo Astrios, Beckman Coulter) with a 100 μm nozzle at a speed of around 5000 cells/sec. Cell analysis was performed on either one of the flow cytometers (Attune NxT, Thermo Fisher; Cytek AURORA, Aurora).

### Single-cell library construction and sequencing

Sorted CD41^+^ FSC^high^ single cells from four mice of a control MK group and an MK group from mice three days upon L. monocytogenes infection each were processed through the Chromium Single Cell Platform using the Chromium Single Cell 3’ Library and Gel Bead Kit v3 (PN-1000075, 10x Genomics) and the Chromium Single Cell B Chip Kit (PN- 1000074, 10x Genomics) as the manufacturer’s protocol. In brief, over 7000 cells were loaded onto the Chromium instrument to generate single-cell barcoded droplets. Cells were lysed and barcoded reverse transcription of RNA occurred. The library was prepared by following amplification, fragmentation, adaptor, and index attachment then sequenced on an Illumina NovaSeq platform.

### scRNA-seq processing

The scRNA-seq reads were aligned to the mm10 reference genomes, and unique molecular identifier (UMI) counts were obtained by Cell Ranger 3.0.2. Normalization, dimensionality reduction, and clustering were performed with the Seurat 3.0 R package (Butler et al., 2018). For the control and Listeria (L.) monocytogenes infection group, we loaded one 10x Genomics well each and detected 5663 and 5948 cells that passed the Cell Ranger pipeline, respectively. To ruled out low quality cells, cells with >12% mitochondrial content or <200 detected genes were excluded with Seurat function subset (percent.mt<12 & nFeature_RNA>200). We ruled out doublets with default parameters of DoubletDecon R package, and 54 control cells and 939 L. monocytogenes infected cells were excluded. Following the standard procedure in Seurat’s pipeline, we identified 3272 MKs from control mice (1712 MKs) and mice with L. monocytogenes infection (1560 MKs) (3897 and 3449 immune cells were discarded respectively) in combination with MetaNeighbor method. Preprocessed dataset normalization was performed by dividing the UMI counts per gene by the total UMI counts in the corresponding cell and log-transforming before scaling and centering. SCT normalization was performed with the script: object <- SCTransform(object, vars.to.regress = “percent.mt”, verbose = FALSE). Signature genes of each cluster were obtained using the Seurat function FindMarkers with Wilcox test with fold change > 1.5 and p value < 0.05 after clustering. Heatmaps, individual UMAP plots, and violin plots were generated by the Seurat functions in conjunction with ggplot2 and pheatmap R packages.

Similarities and UMAP projection between our scRNA-seq data and published datasets GSE152574 (Yeung et al., 2020), GSE158358 (Pariser et al., 2021), GSE137540 (Xie et al., 2020), GSE128074 (Hamey et al., 2021), or GSE132042 (Almanzar et al., 2020) were conducted by MetaNeighbor R package (Crow et al., 2018), iMAP.py and Symphony R package (Kang et al., 2021). iMAP integration was performed using the default parameters except n_top_genes=2000, min_genes=0, min_cells=0, and n_epochs=100 before doing dimensionality reduction using Uniform Manifold Approximation and Projection method (UMAP, n_neighbors=30, n_pca=30). Radar charts were generated with JavaScript written by Nadieh Bremer (VisualCinnamon.com). Euclidean distances denote the distances between the centroid of each cluster.

Correlations were calculated based on normalized RNA values, with the function cor and the parameter ‘method= “spearman”’. Multiple testing correction using the function cor.test with the parameter “method = “spearman” and it was applied for Cxcr4 expression correlations. We calculated the similarities between MK1 to 5 with the published MK, immune cell, and myeloid progenitor datasets (Almanzar et al., 2020; Hamey et al., 2021; Pariser et al., 2021; Xie et al., 2020; Yeung et al., 2020) using scmap R package (Kiselev et al., 2018). Default parameters and 1 000 features were used and threshold > 0 was set. Cell-type matches are selected based on the highest value of similarities and the second- highest value which is not 0.01 less than the highest value across all reference cell types.

Cytokine, inflammatory, chemokine, and antigen processing and presentation scores were evaluated with the AddModuleScore function of Seurat using genes from KEGG pathway ko04060, cytokine-cytokine receptor interaction; GO:0006954, inflammatory response; chemokine ligands from CellPhoneDB.mouse (Jin et al., 2021) and GO:0019882, antigen processing and presentation.

Interaction analysis of MKs and immune cells were conducted by CellPhoneDB (Efremova et al., 2020) (transformed to human orthologous genes (Davidson et al., 2020)) and CellChat R package (Jin et al., 2021). Only interactions involving cytokines were shown. Gene Ontology (GO) analysis was performed using clusterProfiler R package (Yu et al., 2012) and visualized using enrichplot R package (Yu, 2019).

Gene set enrichment analysis (GSEA) was performed using gsea R package (Subramanian et al., 2005) and visualized using enrichplot R package. Gene lists were pre- ranked by the fold change values of the differential expression analysis using Seurat function FindMarkers. Gene sets for GSEA were obtained from GO database (GO:0002367, cytokine production involved in immune response; GO:0006954, inflammatory response; GO:0008009, chemokine activity; GO:0022409, positive regulation of cell-cell adhesion; GO:0002275, myeloid cell activation involved in immune response).

Gene set variation analysis (GSVA) was performed using GSVA R package (Hanzelmann et al., 2013). GSVA was performed to calculate GSVA score of indicated pathway genes in single cell datasets with the whole protein encoding genes after log normalization of expression values. Gene sets for GSVA were obtained from GO database (GO:0022409, positive regulation of cell-cell adhesion; GO:0002275, myeloid cell activation involved in immune response; GO:0002367, cytokine production involved in immune response; GO:0007596, blood coagulation; GO:0019882, antigen processing and presentation; GO:0034340: response to type I interferon; GO:0034341: response to interferon-gamma; GO:0045088, regulation of innate immune response; GO:0042742, defense response to bacterium; GO:0002819, regulation of adaptive immune response; GO:1903708, positive regulation of hemopoiesis).

### Lung cells preparation for flow cytometry

Lungs were removed and digested as described with minor modifications (Lefrancais et al., 2017). In brief, removed lungs were placed in 1.5 ml tubes, minced with scissors, and digested with 1 ml digestion buffer (HBSS with 1mg/ml collagenase D, 0.1 mg/ml DNase I, 25 mM HEPES, 2 mM L-glutamine, and 2% FBS) for 30 min at 37℃ before filtration through a 100-μm cell strainer and red blood cell lysis for 5 min. Samples were then filtered through 70-μm strainers and resuspended for subsequent surface marker staining for flow cytometry.

### Megakaryocyte ablation induction

Pf4-cre mice were mated with the iDTR line to generate Pf4-cre; iDTR mice. Diphtheria toxin (DT, Sigma-Aldrich) was injected intraperitoneally every day at a dose of 40 ng g^−1^ bodyweight into Pf4-cre^+^; iDTR^+/–^ mice and their cre negative counterparts to induce megakaryocyte ablation as indicated.

### Cre-ER recombinase induction

Scl-creER mice were mated with the tdTomato line to generate Scl-creER; tdTomato mice. For induction of cre-ER recombinase, Scl-creER, tdTomato^+/–^ mice were injected with tamoxifen intraperitoneally once (2 mg in 0.1 ml corn oil; Sigma-Aldrich).

### BrdU incorporation assay

5-bromo-2-deoxyuridine (BrdU) was administered at a single dose of 125 mg kg^−1^ body mass by intraperitoneal injection. Whole bone marrow cells were collected 12 hours later and incubated with anti-CD41 and anti-CXCR4 for one hour. Cells were washed and then fixed with 4% PFA at 4 °C overnight. Cells were then permeabilized with 0.5% TritonX- 100 for 15 minutes at room temperature and incubated with 1mg ml^−1^ DNase I (Roche) for one hour at 37 °C followed by incubating with anti-BrdU (BU20A, eBioscience) for one hour at room temperature before being analyzed.

### Annexin V binding assay

For Annexin V binding assay, bone marrow cells were incubated with cell surface markers for one hour at 4 °C and then washed with PBS before being resuspended with Annexin V binding buffer (Biolegend). Cells were then incubated with FITC Annexin V (Biolegend) for 15 minutes at room temperature in dark, and then 300 μl Annexin V binding buffer was added to each tube. Cells were analyzed by a flow cytometer.

### Immunostaining

Immunostaining of frozen sections was performed as described (Jiang et al., 2018). For bone sections, mice were perfused with PBS and 4% paraformaldehyde (PFA). Then the bones were fixed with 4% PFA for 24 hours, decalcified with 0.5 M EDTA for 2 days, and gradient dehydrated by 15% and 30% sucrose for another 2 days. The thick of sections was 30 μm. We used CD41 (MWReg30; eBioscience; 1:200), Endomucin (R&D; 1:100), CD150 (TC15-12F12.2; Biolegend; 1:100), CD48 (HM48-1; Biolegend; 1:100), CXCR4 (2B11, eBioscience; 1:100) antibodies, and lineage panel (Biolegend; cat #133307; 1:50). Secondary staining was done with donkey anti–goat AlexaFluor 488 (Invitrogen; 1:1000). For the liver and spleen from Pf4-cre^+^; mT/mG^+/–^ mice, and lung from Pf4-cre^+^; tdTomato^+/–^ mice, we used DAPI (Thermo Fisher; 0.5 μg ml^−1^) to stain the frozen sections. For phagocytosis analysis, F4/80 (BM8, eBioscience; 1:100), CD11b (M1/70; Invitrogen; 1:100), CD41 (MWReg30; Thermo Fisher; 1:200) and DAPI was used. For sorted MKs, we used CXCR4 (2B11, eBioscience; 1:100), TNFα (R023, Sino Biological; 1:100) and IL-6 (MP5-20F3, Biolegend; 1:100) antibody. Secondary staining was performed with donkey anti–rabbit AlexaFluor 488 (Invitrogen; 1:1000). Confocal images were obtained using a spinning-disk confocal microscope (Dragonfly, Andor) and analyzed using Imaris 9.0 software (Oxford Instruments). Three-Dimension plots were generated using Matplotlib (Hunter, 2007).

### Quantitative real-time (qRT-) PCR

For RT-qPCR, MKs were dissociated in Trizol (Magen), and RNA was extracted following the manufacture’s instruction. RNA was reverse transcribed into cDNA using the TransCript All-in-One First-Strand cDNA Synthesis kit (Transgene). Quantitative PCR was performed using a Bio-Rad CFX 96 touch. The primers for Pf4 were 5’- GGGATCCATCTTAAGCACATCAC-3’ (forward) and 5’-CCATTCTTCAGGGTGGCTATG-3’ (reverse). The primers for Vwf were 5’- CTTCTGTACGCCTCAGCTATG-3’ (forward) and 5’- GCCGTTGTAATTCCCACACAAG-3’ (reverse). The primers for Mpl were 5’-AACCCGGTATGTGTGCCAG-3’ (forward) and 5’-AGTTCATGCCTCAGGAAGTCA-3’(reverse). The primers for Cxcl12 were 5’-AGGTTCTTATTTCACGGCTTGT-3’ (forward) and 5’-TGGGTGCTGAGACCTTTGAT-3’ (reverse). The primers for Gapdh were 5’- AGGTCGGTGTGAACGGATTTG-3’ (forward) and 5’-GGGGTCGTTGATGGCAACA-3’ (reverse). Gapdh was used as the reference gene for qRT-PCR analysis.

### Transwell migration

Transmigration assays were performed on a transwell with a pore size of 5 μm (Biofil). CXCR4^low^ MKs or CXCR4^high^ MKs from bone marrow were sorted (5000 cells per well) from control mice and added to the lower chamber with 600 μl IMDM (Thermo Fisher) plus 10% FBS (Gibco). Peripheral blood cells were collected as described in the “Flow cytometry and cell sorting” section. 6 ×10^5^ peripheral blood cells were resuspended in 100 μl RPMI 1640 (Gibco) plus 10% FBS and added to the upper insert to continue for two- hour incubation at 37 °C, 5% CO2. Cells in the lower chamber were harvested, washed with PBS once, and resuspended with 100 μl PBS for staining and FACS counting.

### Phagocytosis

Bone marrow-derived macrophages (BMDM) from C57BL/6 mice at 6-8 weeks of age were differentiated from bone marrow precursors with minor modifications (Minutti et al., 2019). In brief, bone marrow cells were isolated and propagated for seven days in DMEM without sodium pyruvate or HEPES (Gibco), containing 20% FBS (Gibco), 30% supernatants of L929 conditioned media, and 1% Pen/Strep (Hyclone) at 37 °C. Macrophage phagocytosis assays were performed on a transwell plate with a pore size of 5 μm (Biofil) as described with modifications (Sharif et al., 2014). Briefly, attached cellswere replated into 24-well plates, 5×10^4^ cells per well, on glass coverslips for 24-hour culture. Then 5 000 sorted CXCR4^low^ MKs or CXCR4^high^ MKs were added in the upper inserts and placed onto macrophages chambers for additional 16 hours incubation without or with 2 μg ml^−1^ TNFα neutralizing antibody (R023, Sino Biological; 1:100) or 2 μg ml^−1^ IL-6 neutralizing antibody (MP5-20F3, Biolegend) at 37 °C, 5% CO2. The upper inserts were discarded and macrophages were washed with PBS without antibiotics and incubated with 10^5^ GFP-labeled E. coli for two hours at 37 °C, 5% CO2. Cells were washed three times with PBS and incubated with DMEM without sodium pyruvate or HEPES (Gibco) with gentamycin (50 μg ml^−1^) for 30 minutes at 37 °C, 5% CO2 to remove adherent bacteria. Cells were then fixed by cold methanol for 15 minutes and blocked with 10% BSA overnight, followed by incubation with F4/80 (BM8, eBioscience; 1:100) for two hours at room temperature before being quantified using a spinning disk confocal microscope (Dragonfly, Andor).

For neutrophil phagocytosis, CD11b^+^ Gr1^+^ Ly6c^−^ neutrophils were sorted from the spleen and propagated in RPMI 1640 (Gibco) containing 10% FBS. Neutrophil phagocytosis was performed as described in macrophage phagocytosis assay, except cells were sedimented for 30 minutes and fixed on glass coverslips after incubated with GFP-E. coli and gentamycin. The capacity of phagocytosis was evaluated by fluorescence intensity of GFP-E. coli. within macrophages and neutrophils.

### Bone marrow live imaging

Pf4-cre^+^; tdTomato^+/–^ mice were infected with L. monocytogenes for 24 hours. FITC- Dextran (average mol wt 2000000, Sigma-Aldrich) was injected intravenously at a dose of 1.25 mg per mouse before being sacrificed. Bone marrow was flushed integrally and fixed onto a glass slide in a chamber, rinsed with RPMI 1640 (Gibco), and covered slightly with a coverslip. Confocal images were obtained every minute on the spinning-disk confocal microscope (Dragonfly, Andor) and analyzed using Imaris 9.0 software (Oxford Instruments).

### In vitro MK culture, MK size, and proplatelet formation measurement

MKs were sorted using a cell sorter (MoFlo Astrios, Beckman Coulter) and cultured in 24- well plates in SFEM (Stem Cell Technologies) plus 100 ng ml^−1^ mTPO (Novoprotein) and 1% Pen/Strep (Hyclone), and incubated at 37 °C, 5% CO2 for four days. Images were taken by a Nikon Ts2R microscope equipped with a Nikon DS-Ri2 camera. Cell size and proplatelet formation were measured on day three or day five post-cultured, respectively, using Nikon NIS-Elements BR.

### Bone marrow transfer experiments

Pf4-cre mice were mated with the tdTomato line to generate Pf4-cre^+^; tdTomato^+/–^ mice. tdTomato^+^ MKs were isolated from Pf4-cre^+^; tdTomato^+/–^ mice. 6-8-week-old recipient mice were pre-treated with PBS or 2500 CFUs of L. monocytogenes as previously described one day before cell perfusion. 1 × 10^5^ tdTomato^+^ MKs were sorted and intravenously injected into control or L. monocytogenes infected mice. tdTomato^+^ MKs were detected in lungs with immunostaining two days after cell perfusion.

mtdTomato^+^ bone marrow cells were isolated from Pf4-cre^−^; mTmG^+/–^ mice. 1 ×10^6^ mtdTomato^+^ bone marrow cells were intravenously injected into control or one-day-L. monocytogenes infected mice. mtdTomato^+^ MKs were detected in bone marrow, liver, and spleen two days after cell perfusion.

For in vivo CXCR4^high^ MK function assay in MK ablation mice, DT was intraperitoneally injected every day for five days. On the second and fourth days, 2 × 10^5^sorted wild type CXCR4^high^ MKs or CXCR4^low^ MKs were intravenously injected into indicated groups. PBS or 2500 CFUs of L. monocytogenes as previously described were injected intravenously on the third day. Spleen and liver were harvested three days after infection to determine the bacterial burdens as described.

### T cell reactivation in vitro

Splenocytes (1 × 10^6^ cells well^−1^) from control or MK ablated mice after seven days L.m.- OVA infection were re-stimulated for four hours in vitro with OVA peptide (10 μM) in the presence of Brefeldin-A (BFA, 10 μg ml^−1^). Activated T cells were then analyzed by a flow cytometer.

### Computational modeling of random myeloid cell localization

We have performed randomized simulations as in previous reports (Bruns et al., 2014; Jiang et al., 2018) in Python. Images of a 400 μm × 400 μm bone marrow region with CXCR4^high^ and CXCR4^low^ MKs, in which background staining was removed, were used to generate MKs onto which 200 myeloid cells were randomly placed, consistent with an average density of 200 myeloid cells per field. Each simulated run placed 200 random myeloid cells (mean diameter 5 μm) was repeated 500 times. The shortest Euclidean distance was calculated for each myeloid cell to CXCR4^high^ or CXCR4^low^ MKs. Random and observed distance distributions were compared using the modified nonparametric two-dimensional (2D) KS test as described (Bruns et al., 2014; Jiang et al., 2018).

### Statistical analyses

Data are presented as means ± s.e.m except for phagocytosis assays and MK size measurement, which are presented as means ± first and third quartiles. For phagocytosis assay and MK size measurement, data were analyzed by a one-dimensional KS test.

Differences were considered statistically significant if p < 0.05. For the comparison of three-dimensional distances, a two-dimensional KS test was used. The difference was considered statistically significant if p < 0.05. For multiple comparisons analysis, data were analyzed by repeated-measures one-way analysis of variance (ANOVA) followed by Dunnett’s test. Differences were considered statistically significant if p < 0.05. ǂ P<0.05, ǂǂ P<0.01, ǂǂǂ P<0.001, n.s., not significant. For other experiments except for scRNA-seq analysis, data were analyzed by a two-tailed Student’s t-test. Differences were considered statistically significant if p < 0.05. * p < 0.05, ** p < 0.01, *** p < 0.001, n.s., not significant.

### Data availability

The scRNA-seq data generated in this study are deposited in GEO (GSE168224, https://www.ncbi.nlm.nih.gov/geo/query/acc.cgi?acc=GSE168224). The code used in the study can be accessed at GitHub (https://github.com/JYCathyXie/MK_infection).

## Acknowledgements

We thank the National Key Research and Development Program of China (2017YFA0103403, 2018YFA0107200), NSFC (81822001, 81900101, 81800164), The Key Research and Development Program of Guangdong Province (2019B020234002), Shenzhen Foundation of Science and Technology (JCYJ20170818103626421), China Postdoctoral Science Foundation (2021M693614), Guangdong Innovative and Entrepreneurial Research Team Program (2016ZT06S029, 2019ZT08Y485), Sanming Project of Medicine in Shenzhen (No.SZSM201911004) for generous support.

## Author contributions

J.W., J.X., D.W., X.H. designed and performed most of the experiments, analyzed the data, and generated figures. D.W. contributed to histology and live imaging. M.C. contributed to MK ablation experiments. L.J. and G.S. contributed to scientific discussion and manuscript preparation. M.Z. supervised the project and wrote the manuscript.

## Competing interests

The authors declare no competing interests.

## Notes

https://www.ncbi.nlm.nih.gov/geo/query/acc.cgi?acc=GSE168224

### Competing Interest Statement

The authors have declared no competing interest.

