## Supplemental figures for "Megakaryocyte derived immune-stimulating cells regulate host-defense immunity against bacterial pathogens"

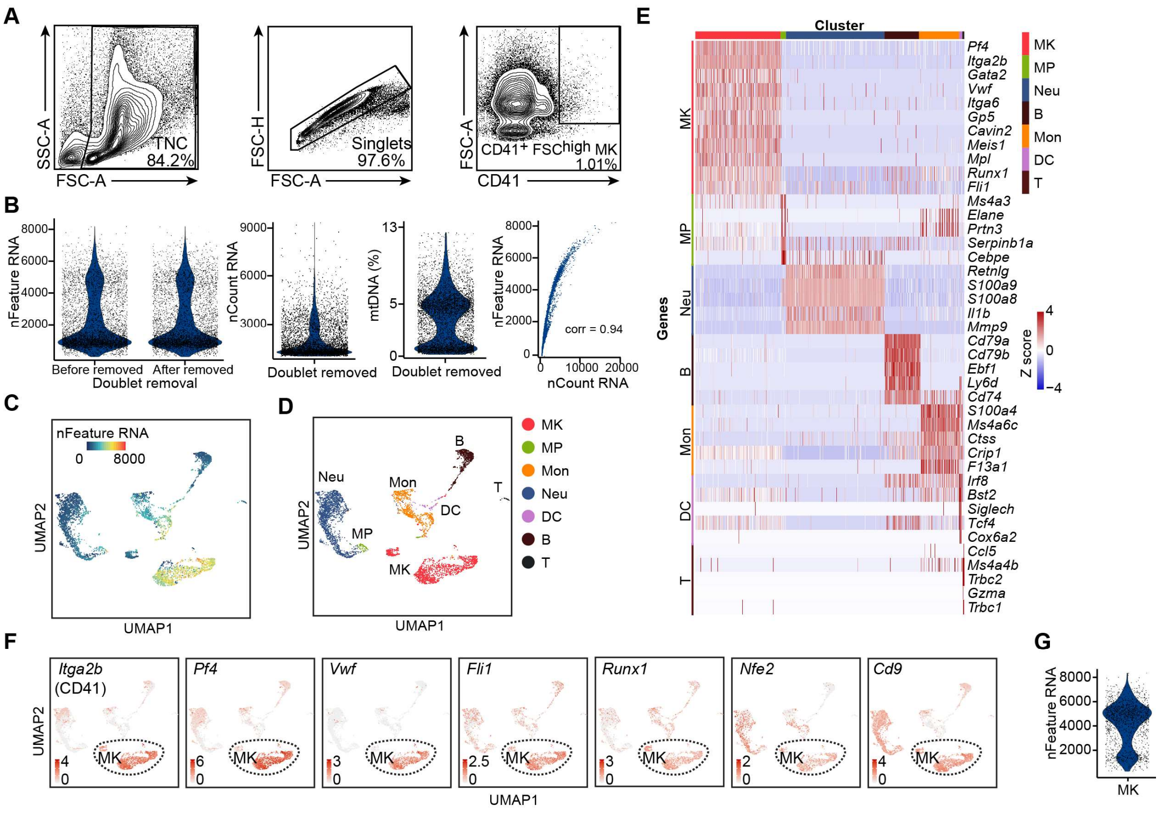


**Figure 1-figure supplement 1. Cell isolation, quality control and annotation of scRNA-seq data.**

(**A**) Flow cytometry gating for isolation of MKs (CD41$^{+}$ FSC$^{high}$) in bone marrow for scRNA-seq. (**B**) Quality control of scRNA-seq data. Violin plots showing the number of unique genes (gene number) before and after removing doublets, number of total unique molecular identifiers (UMI counts) and percentage of mitochondrial transcripts in single cells after removing doublets. Scatter plot showing the correlation between UMI counts and gene numbers. corr indicates Pearson correlation coefficient. (**C-D**) UMAP of 5368 bone marrow cells, colored by gene numbers (**C**) and by cluster identity indicated on the right (**D**). MK, megakaryocytes; Neu, neutrophils; MΦ, macrophages; MP, myeloid progenitors; DC, dendritic cells; Mon, monocytes; B, B cells; T, T cells. (**E**) Heatmap of row-scaled signature gene expression in each cell type (top, color-coded by subpopulations). Columns denote cells; rows denote genes. Z score, row-scaled expression of the signature genes in each subpopulation. (**F**) Feature plots showing selected gene expression. (**G**) Violin plots showing the number of unique genes (gene number) of 1712 MKs.

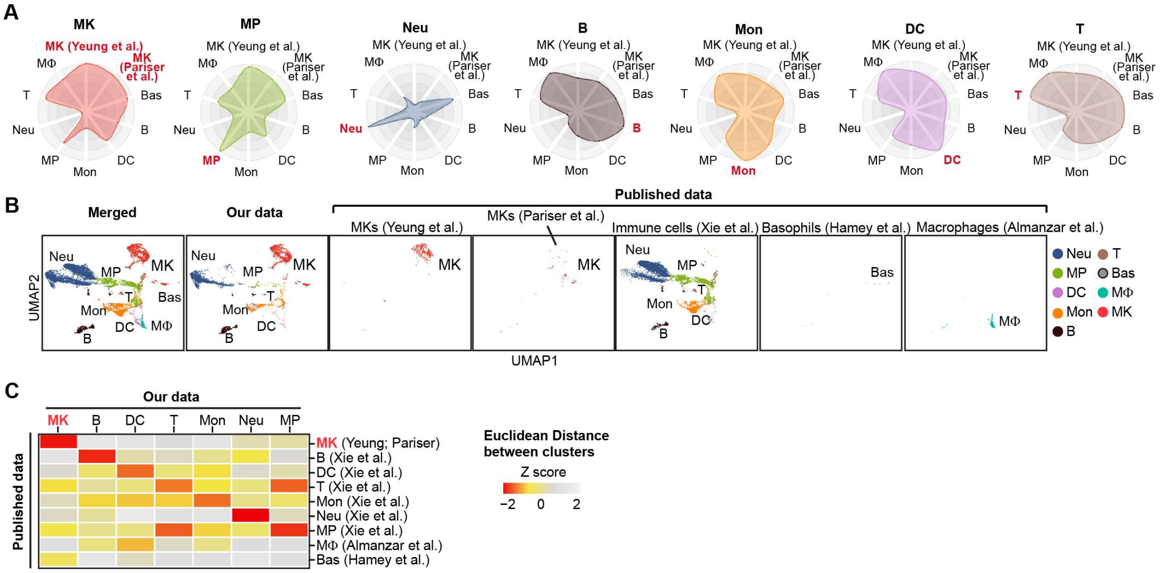

**Figure 1-figure supplement 2. Cell type identification by alignment with published scRNA-seq data.**

(**A**) Radar charts showing cell similarities of our single cell dataset with the published bone marrow MK single cells (Pariser et al., 2021; Yeung et al., 2020), bone marrow immune cells and myeloid progenitor single cells (Almanzar et al., 2020; Hamey et al., 2021; Xie et al., 2020) using MetaNeighbor R package. (**B**) Comparison of our bone marrow scRNA-seq data with published bone marrow MK (Pariser *et al.*, 2021; Yeung *et al.*, 2020) and immune cell (Almanzar *et al.*, 2020; Hamey *et al.*, 2021; Xie *et al.*, 2020) datasets. All cells were integrated by iMAP.py and projected on UMAP. (**C**) Column-scaled Euclidean distances between the centroid of each cluster. Z score, column-scaled Euclidean distance. MK, megakaryocytes; Neu, neutrophils; MΦ, macrophages; MP, myeloid progenitors; DC, dendritic cells; Mon, monocytes; B, B cells; T, T cells.

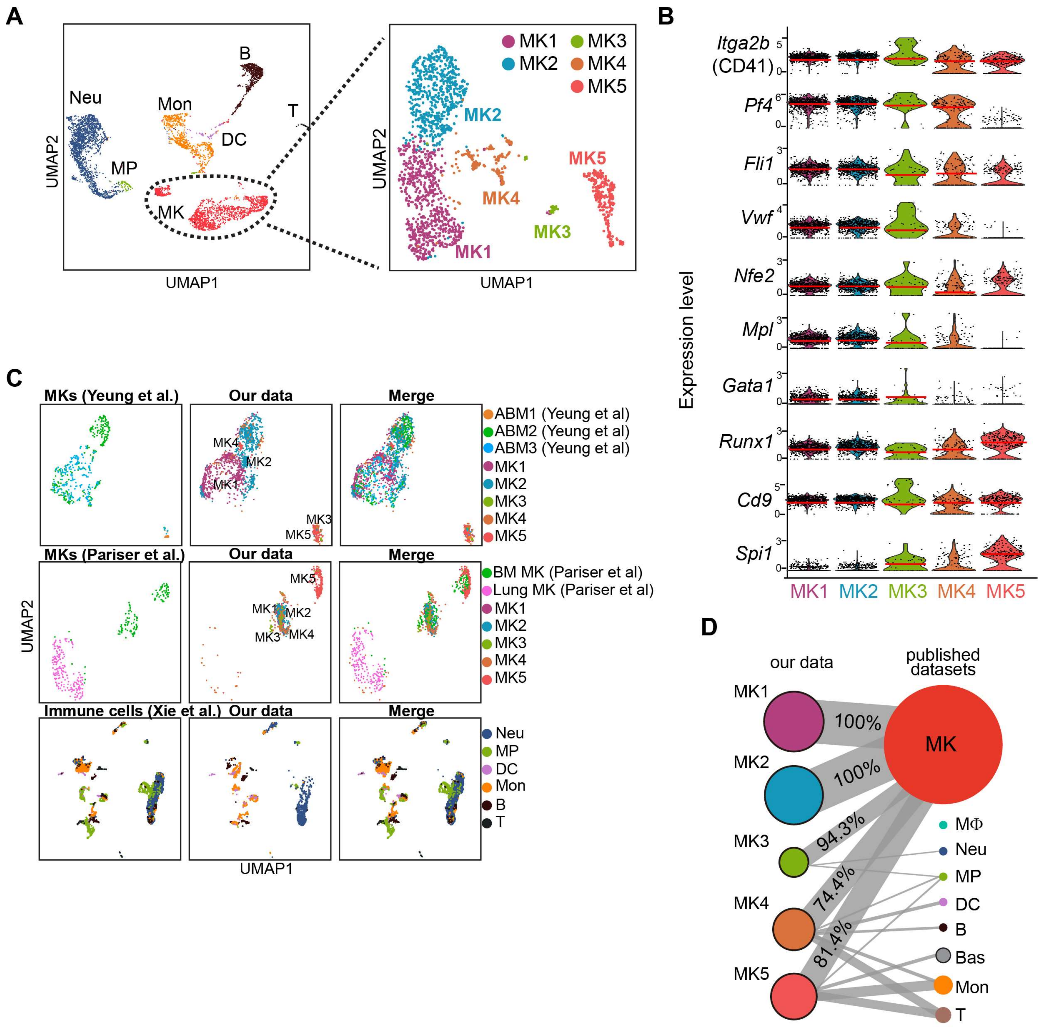


**Figure 1-figure supplement 3. Identification of MK subpopulations.**

(**A**) Re-clustering of 1712 MKs from 5368 cells. (**B**) Violin plots showing MK marker gene expression (Aburima et al., 2021; Bernardes et al., 2020; Kanaji et al., 2005; Liu et al., 2021; Machlus and Italiano, 2019; OZAKI et al., 2005; Sun et al., 2021; Yeung *et al.*, 2020) of MK1 to 5. Red lines indicate the median gene expression. (**C**) Projection of our datasets on reported MK (Pariser *et al.*, 2021; Yeung *et al.*, 2020) and immune cell (Hamey *et al.*, 2021; Xie *et al.*, 2020) scRNA-seq datasets by Symphony R package. MK, megakaryocytes; Neu, neutrophils; MΦ, macrophages; MP, myeloid progenitors; DC, dendritic cells; Mon, monocytes; Bas, basophils; B, B cells; T, T cells; ABM, adult bone marrow. (**D**) MKs (MK1 to 5) and cell types projection based on similarities of our single cell transcriptional profiles and published MK (Pariser *et al.*, 2021; Yeung *et al.*, 2020) and immune cell datasets (Hamey *et al.*, 2021; Xie *et al.*, 2020) by scmap R package.

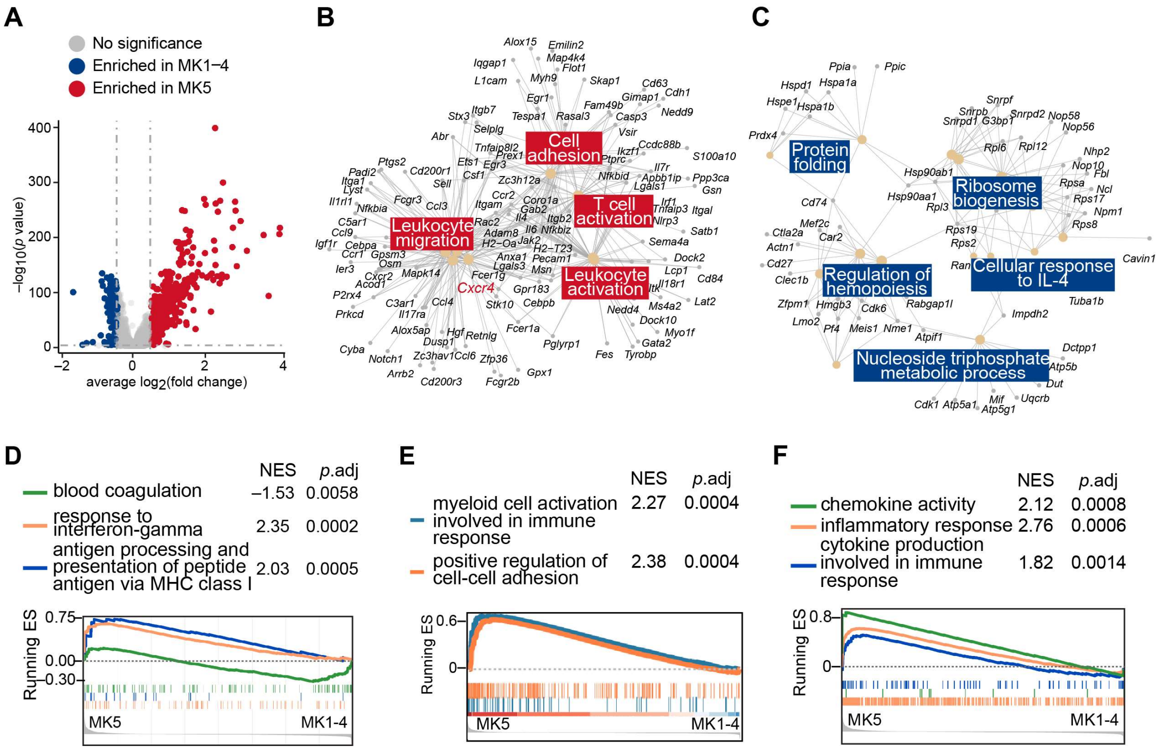


**Figure 1-figure supplement 4. Enriched genes in MK1 to 4 and MK5.**

(**A**) Volcano plot showing MK5 and MK1 to 4 enriched genes (|fold change| > 1.4, $p$ value < 0.05). (**B-C**) Gene Ontology (GO) analysis showing MK5 enriched immune pathway genes and MK1 to 4 mainly enriched hemopoiesis and RNA processing pathway genes. (**D-F**) Gene set enrichment analysis (GSEA) evaluated enrichment of selected pathways in MK5 cells compared to other MK subpopulations (MK1 to 4).

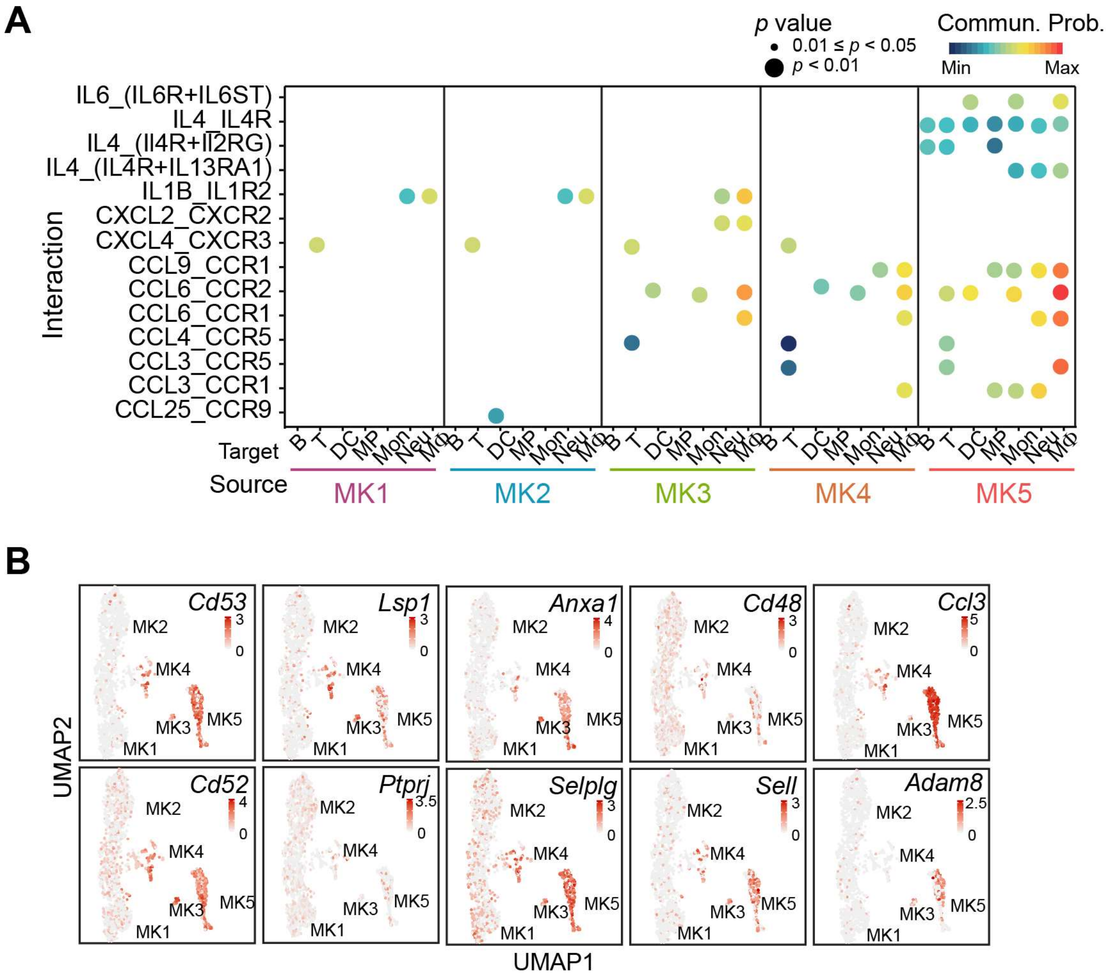


**Figure 1-figure supplement 5. MK5 interacts with immune cells and express signature genes of immune MKs.**

(**A**) Dotplots of significant cytokine ligand (source) -receptor (target) interactions between MKs and immune cells discovered using CellChat. Color indicates communication probabilities, and bubble size indicates *p* value of the ligand-receptor pairs between MK subpopulations (source) and immune cells (target). (**B**) Selected signature gene expression in MK subpopulations.

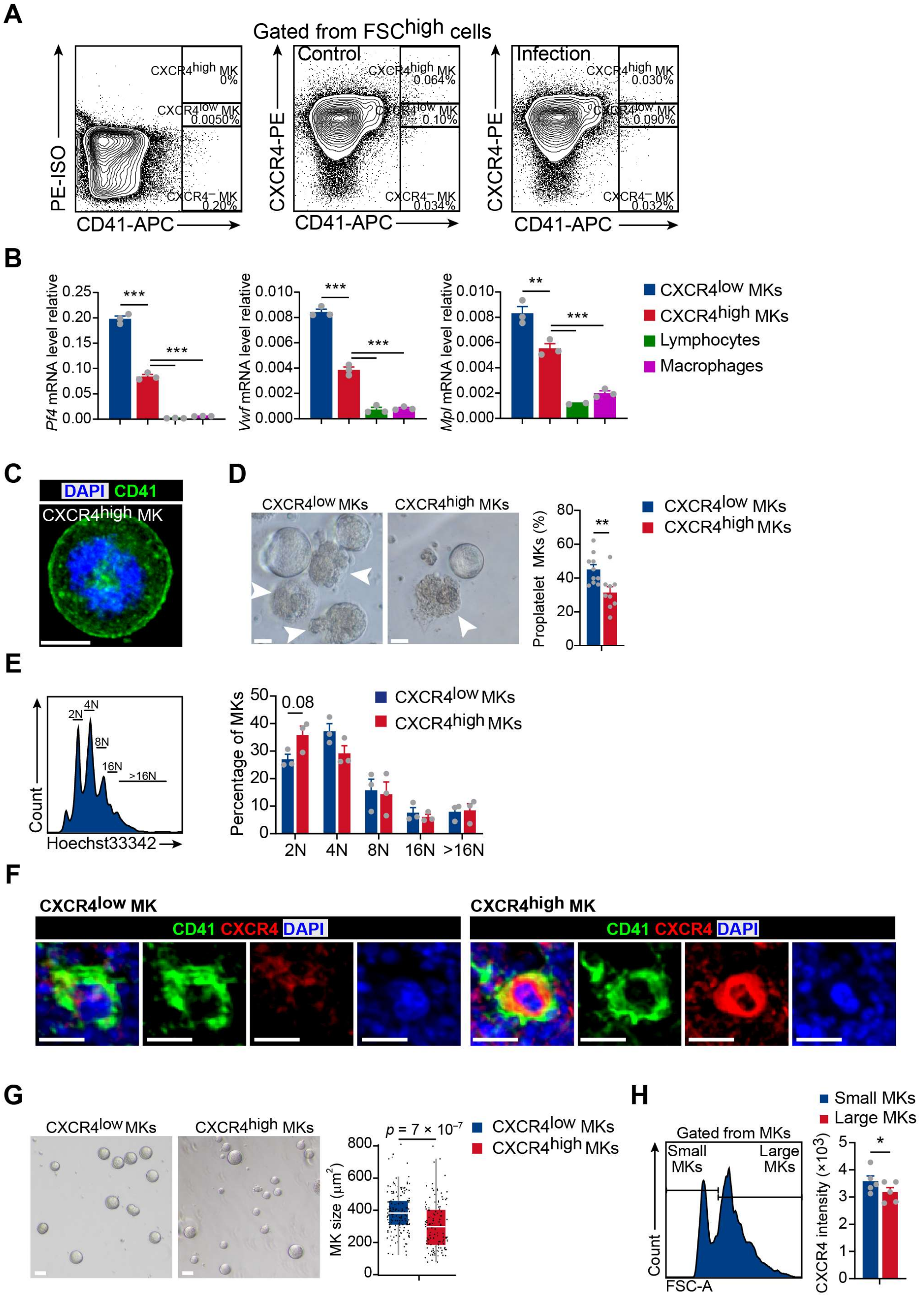


**Figure 1-figure supplement 6. Polyploidy, platelet generation ability and cell size of CXCR4$^{low}$ and CXCR4$^{high}$ MKs.**

(**A**) Representative flow cytometry plots of gating strategy of CD41 and CXCR4 in the bone marrow of control mice and mice three days after *L. monocytogenes* infection. (**B**)

Relative expression of *Pf4*, *Vwf* and *Mpl* in CXCR4$^{high}$ and CXCR4$^{low}$ MKs by RT-qPCR. (**C**) Representative immunofluorescent staining image showing membranous CD41 staining typical of sorted bone marrow CXCR4$^{high}$ MKs. (**D**) Sorted CXCR4$^{low}$ and CXCR4$^{high}$ MKs produced proplatelets *in vitro* on day five post cultured under 100 ng ml$^{-1}$ TPO. White arrowheads indicate proplatelet formation. (**E**) Polyploidy distribution of CXCR4$^{low}$ MKs and CXCR4$^{high}$ MKs (right). (**F**) Representative immunofluorescent staining images showing CD41 and CXCR4 expression of CXCR4$^{low}$ and CXCR4$^{high}$ MKs in the bone marrow. CD41, green; CXCR4, red; DAPI, blue. (**G**) Comparison of cell size between CXCR4$^{low}$ MKs and CXCR4$^{high}$ MKs on day three post cultured under 100 ng ml$^{-1}$ Thrombopoietin (TPO). (**H**) Median fluorescence intensity of CXCR4 in small and large sizes of MKs by flow cytometry. Scale bars, 20μm. Data represent mean ± s.e.m in (**B**, **D**, **E**, **H**) or mean ± first and third quartiles in (**G**). Two-sample KS test was performed to assess statistical significance in (**G**). Two-tailed Student's *t*-test was performed to assess statistical significance in (**B**, **D**, **E**, **H**). * $p$ <0.05, ** $p$ <0.01, *** $p$ <0.001.

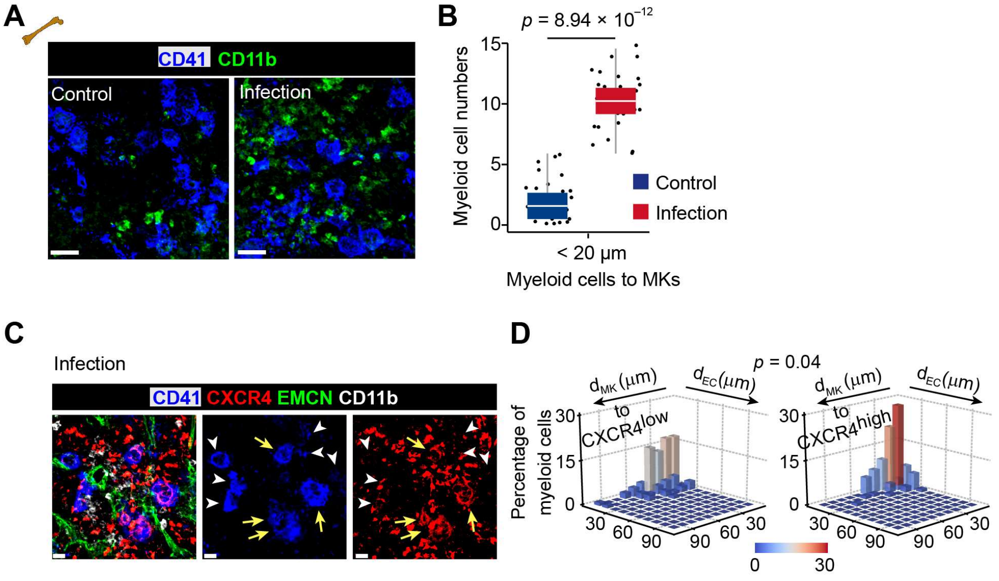


**Figure 2-figure supplement 1. *L. monocytogenes* promote myelopoiesis and the association of myeloid cells and the CXCR4$^{high}$ MK-blood vessel intersection.**

(**A-B**) Representative immunofluorescent staining image (**A**) and statistical analysis (**B**) showing association of MKs (blue) and myeloid cells (green) in the bone marrow of control mice and mice three days after *L. monocytogenes* infection ($n$ = 30 control and infected MKs). CD41, blue; CD11b, green. (**C-D**) Representative images of MKs (blue), CXCR4 (red), vascular endothelial cells (green) and myeloid cells (white) (**C**), and statistical analysis (**D**) in bone marrow showing the distribution of myeloid cells to CXCR4$^{low}$ or CXCR4$^{high}$ MKs and vascular cells three days after *L. monocytogenes* infection. CD41, blue; CXCR4, red; EMCN, green; CD11b, white. Yellow arrows indicate CXCR4$^{high}$ MKs and white arrowheads indicate CXCR4$^{low}$ MKs (left) and two-dimensional probability distribution of distances from myeloid cells to CXCR4$^{low}$ or CXCR4$^{high}$ MKs and vascular cells ($n$ = 104 CD11b$^{+}$ cells; $p$ = 0.04 by 2D KS test). Scale bars, 20μm. Data represent mean ± first and third quartiles in (**B**). Two-sample KS test was performed to assess statistical significance in (**B**). Two-dimensional-two-sample KS test was performed to assess statistical significance in (**D**).

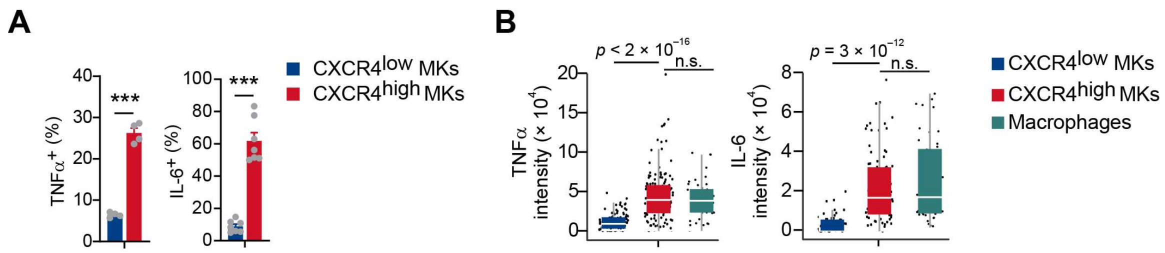


**Figure 2-figure supplement 2. TNFα and IL-6 expression in CXCR4$^{low}$ and CXCR4$^{high}$ MKs.**

(**A-B**) TNFα and IL-6 protein levels in CXCR4$^{low}$ and CXCR4$^{high}$ MKs were shown by flow cytometry (**A**) and immunofluorescent staining (**B**) using sorted MKs, comparing to their levels in sorted macrophages upon *L. monocytogenes* infection (**B**). Data represent mean ± s.e.m in (**A**) or mean ± first and third quartiles in (**B**). Two-sample KS test was performed to assess statistical significance in (**B**). Two-tailed Student's *t*-test was performed to assess statistical significance in (**A**). *** $p$ <0.001.

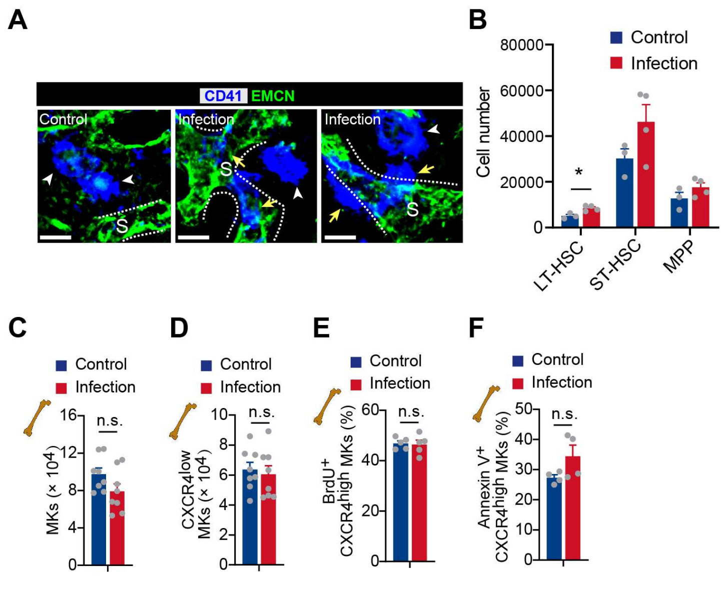


**Figure 4-figure supplement 1. The effects of association of MKs and blood vessels, HSC activation, and MK numbers in bone marrow upon bacterial infection.**

(**A**) Bone marrow MKs (blue) and sinusoids (green) in control mice and mice three days after *L. monocytogenes* infection. CD41, blue; EMCN, green. Yellow arrows indicate MKs egressed into sinusoids. “S” indicates sinusoid and dashed lines demarcate the border of sinusoids. (**B**) HSC and progenitor cell number in the bone marrow of control mice and mice three days after *L. monocytogenes* infection. LT, long term; ST, short term; MPP, multipotential progenitor. (**C-D**) MK numbers (**C**) and CXCR4$^{low}$ MK numbers (**D**) in the bone marrow of control mice and mice three days after *L. monocytogenes* infection ($n$ = 8 mice). (**E-F**) Fraction of BrdU$^{+}$ (**E**) and Annexin V$^{+}$ (**F**) MKs in CXCR4$^{high}$ MKs in bone marrow from control mice or mice three days after *L. monocytogenes* infection ($n$ = 5 mice). Scale bars, 20µm. Data represent mean ± s.e.m. Two-tailed Student’s *t*-test was performed to assess statistical significance. * $p < 0.05$, n.s., not significant.

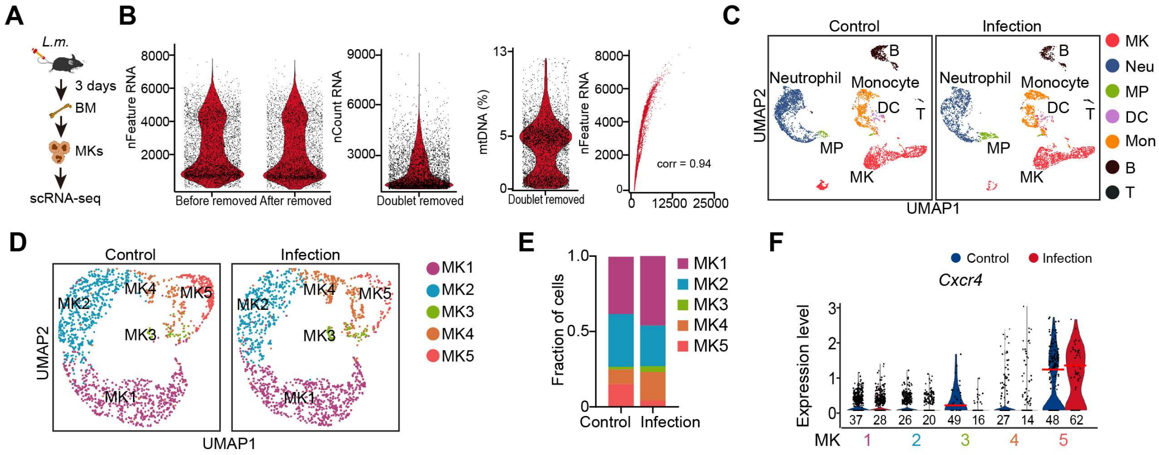


**Figure 4-figure supplement 2. scRNA-seq of MKs from mice upon bacterial infection.**

(**A**) Schematic depicting the strategy of scRNA-seq using bone marrow MKs from mice three days after *L. monocytogenes* infection. (**B**) Violin plots showing the number of unique genes (gene number) before and after removing doublets, number of total unique molecular identifiers (UMI counts) and percentage of mitochondrial transcripts in single cells after removing doublets. Scatter plot showing the correlation between UMI counts and gene numbers. corr indicates Pearson correlation coefficient. (**C**) UMAP of the combined 5 368 cells from control bone marrow and 4 276 cells from *L. monocytogenes* infection bone marrow, colored by cell types. Neu, neutrophil; MP, myeloid progenitor; Mon, monocytes; B, B cells; T, T cells. (**D**) UMAP of the combined 1 712 control MKs and 1 560 infection MKs in the bone marrow, colored by clusters. (**E**) Fraction of each MK subpopulation from control MKs or *L. monocytogenes* infection MKs. (**F**) Violin plot showing *Cxcr4* expression in each MK subpopulation of control and infection MKs.

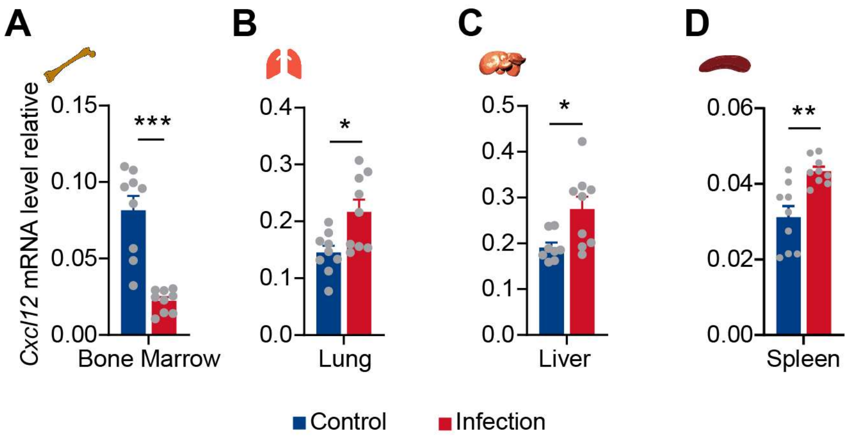


**Figure 4-figure supplement 3.** ***Cxcl12*** **expression upon bacterial infection.**

(**A-D**) Relative expression of *Cxcl12* in the bone marrow (**A**), lung (**B**), liver (**C**), and spleen (**D**) from control mice and mice three days after *L. monocytogenes* infection by RT-qPCR.

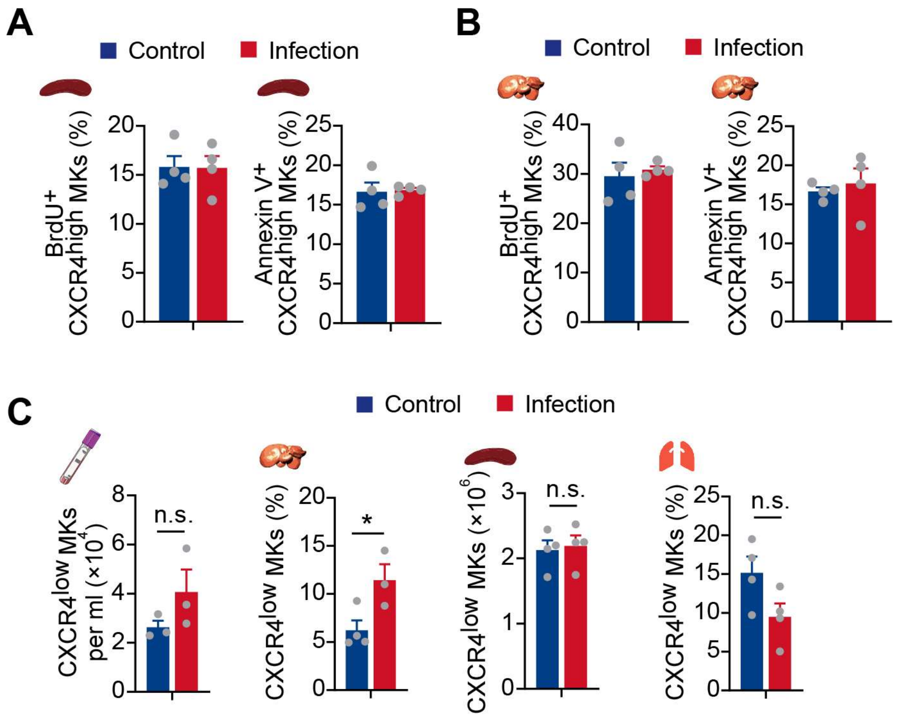


**Figure 4-figure supplement 4. Cell cycle and apoptosis of CXCR4$^{high}$ MKs, and CXCR4$^{low}$ MK numbers in different organs upon bacterial infection.**

(**A-B**) Fraction of BrdU$^{+}$ and Annexin V$^{+}$ MKs in CXCR4$^{high}$ MKs in the liver (**A**) and spleen (**B**) from control mice or mice three days after *L. monocytogenes* infection. (**C**) Quantification of CXCR4$^{low}$ MKs in peripheral blood, liver, spleen, and lung of control mice and mice at three days after *L. monocytogenes* infection. Data represent mean ± s.e.m. Two-tailed Student's *t*-test was performed to assess statistical significance. * $p < 0.05$, n.s., not significant.

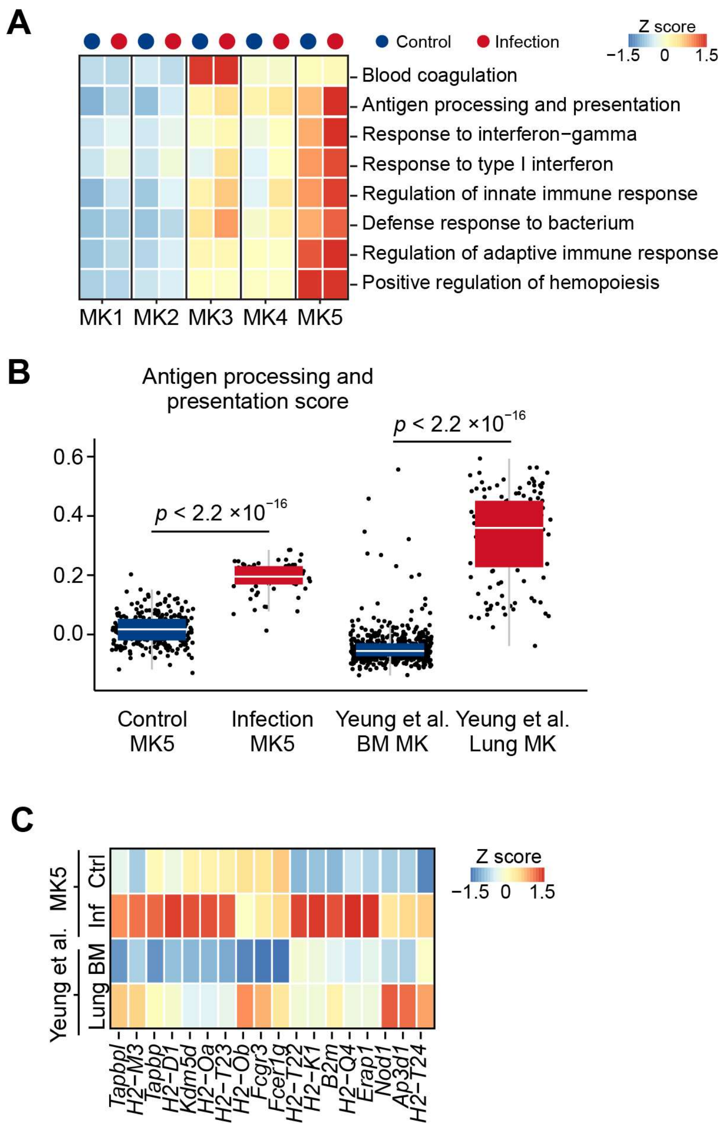


**Figure 4-figure supplement 5. Immune gene expression in bone marrow and lung MKs.**

(**A**) Gene set variation analysis (GSVA) of each MK subpopulation under control or *L. monocytogenes* infection, colored by row-scaled GSVA enrichment scores. (**B**) Antigen processing and presentation score of control MK5, infection MK5, and bone marrow and lung MKs from Yeung et al (Yeung *et al.*, 2020). (**C**) Heatmap showing the row-scaled antigen processing and presentation gene expression of control and infection MK5 cells, comparing with bone marrow MK and lung MK from Yeung et al (Yeung *et al.*, 2020). Data represent mean ± first and third quartiles in (**B**). Two-sample KS test was performed to assess statistical significance in (**B**).

10.1182/bloodadvances.2020002843.
